# RNA-programmable cell type monitoring and manipulation in the human cortex with CellREADR

**DOI:** 10.1101/2024.12.03.626590

**Authors:** Elizabeth A. Matthews, Jeffery B. Russ, Yongjun Qian, Shengli Zhao, Peyton Thompson, Muhib Methani, Matthew L. Vestal, Z. Josh Huang, Derek G. Southwell

**Author notes:** College of Future Technology, Peking-Tsinghua Cener for Life Sciences, IDG/McGovern Institute for Brain Research, Beijing Advanced Center of RNA, Peking University, Peking China. Department of Neurosurgery, Dartmouth Hitchcock Medical Center, Lebanon, NH USA.

## Abstract

Reliable and systematic access to diverse cell types is necessary for understanding the organization, function, and pathophysiology of human neural circuits. Methods for targeting human neural populations are scarce and currently center on identifying transcriptional enhancers and engineering viral capsids. Here we demonstrate the utility of CellREADR, a programmable RNA sensor-effector technology that couples cellular RNA sensing to effector protein translation, for accessing, monitoring, and manipulating specific neuron types in human cortex, *ex vivo*. We designed CellREADRs to target two subpopulations, *CALB2* GABAergic interneurons and *FOXP2* glutamatergic projection neurons, then validated targeting specificity using histological, electrophysiological, and transcriptomic methods. CellREADR expression of channelrhodopsin and GCamp enabled the manipulation and monitoring of these populations in live cortical microcircuits. By demonstrating specific, reliable, and programmable experimental access to human neuronal subpopulations, our results highlight CellREADR’s potential for studying neural circuits and treating brain disorders.

## INTRODUCTION

The human brain has evolved divergent and species-specific properties that underlie our unique cognitive and behavioral capabilities. While approaches such as MRI, EEG, and post-mortem neuroanatomical studies have identified many brain areas, functional networks, and fiber tracts involved in human sensorimotor, cognitive, and emotional functions and their related disorders, brain function fundamentally emerges from neural circuit computations performed by diverse, synaptically interacting cell populations. To better understand the neural underpinnings of human behaviors and neurological conditions, it is thus essential to investigate the fundamental elements of the brain - the diverse human neural cell types - and to dissect their functional features and dynamic interactions at the spatiotemporal resolution of neural circuit operation^1,2^. This requires 1) identifying and categorizing the diverse cell populations that comprise the brain, 2) understanding the input-output connectivities of these cell types, and 3) characterizing their physiological functions in circuit activity. Tools that allow for specific and systematic access to diverse human cell types are essential to these efforts.

In recent years, single-cell RNA sequencing (scRNA-Seq) studies have yielded transcriptomic cell type taxonomies of the rodent, non-human primate (NHP), and human brains^3–12^. In particular, scRNA-seq data have demonstrated conserved as well as divergent features of human cell types, brain areal specializations in cell composition^4,5^, and evidence for genetic adaptations that may confer human-specific neural circuit architectures^3,7,9,13^. Importantly, these transcriptional data also serve as a catalog of genetic access points (i.e., “marker genes”) that may enable the systematic targeting of human cell types for integrated multimodal analyses. What has been lacking, however, are reliable, facile and scalable tools that leverage gene expression to provide experimental access to marker-defined cell populations for human neuroscience^2,14^.

Traditionally, cell type experimental access has relied on germline engineering approaches in model organisms (e.g., transgenic mouse driver lines)^15–18^, which are inapplicable to human research^15–17^. More recently, enhancer-based viral vectors have been advanced as an alternative, cross-species approach to targeting neural cell types^19–22^. However, because enhancers work through complex DNA-protein interactions across complex chromatin landscapes, they often require large-scale efforts to identify and validate^21,23,24^. Furthermore, as enhancer tools are typically validated in mouse models, their generalizability and applicability to human cell types remain to be established. Adeno-associated viral capsid engineering is another potential strategy for cell type access^25^, but capsid tropism at the resolution of specific cell types has yet to be demonstrated for neurons, particularly in human brain tissues^26–30^.

Neurosurgical tissue specimens represent a promising experimental platform for tool development and human neuroscience research. Resected cortical tissues can be maintained *ex vivo* for weeks in organotypic culture preparations, which maintain local tissue cytoarchitecture, cell transcriptomic profiles, morphological features, and neurophysiological activity^31–33^. *Ex vivo* human tissues have been used for a variety of live, semi-chronic research applications, such as the characterization of human disease pathogenesis^34^, testing of emerging therapeutic approaches^35,36^, and validation of novel experimental tools^21,22,37^. They have also been increasingly utilized for morphological and electrophysiological studies of human neurons^38–42^ with, in some cases, subsequent analyses of cell transcriptomic profiles^6,33,43–45^. By enabling researchers to overlay cellular anatomical and physiological data onto transcriptomic datasets, this set of approaches has enabled an initial multi-modal profiling of some human brain cell types. However, previous studies have restricted neuronal sampling based on visible cellular features, such as soma shape and laminar position^6,46,47^, or through viral targeting with broadly active enhancers^19–21^. Altogether a lack of specific and effective methods for accessing molecularly defined cell types has limited the basic and pre-clinical potential of human *ex vivo* tissue preparations.

We recently developed a technology orthogonal to enhancers, CellREADR, which uses a RNA and translation-based method to achieve cell type-specific targeting across diverse organisms^48^. CellREADR - Cell access through RNA sensing by Endogenous ADAR (adenosine deaminase acting on RNA) - is a programmable RNA sensor-effector technology that couples the detection of a cell-defining RNA to the translation of a user-selected effector protein; this coupling is achieved through an ADAR-mediated A-to-I editing mechanism ubiquitous to metazoan cells. Two independent teams have described essentially similar methods, and termed them RADARS^49^ and RADAR^50^. In a prior study we characterized CellREADR sensor and effector properties in human cell lines, demonstrated CellREADR-mediated monitoring and manipulation of neurons in behaving mice, and provided initial proof-of-principle for its application in *ex vivo* human brain tissues^48^. These initial results highlighted CellREADR’s potential as a specific, scalable, and versatile genetic tool for studying human brain cell types and neural circuits.

In this study we demonstrate CellREADR’s utility for accessing, monitoring, and manipulating marker-defined cell types in *ex vivo* human brain. We designed CellREADR constructs to provide experimental access to *CALB2* (calretinin) GABAergic interneurons and *FOXP2* (forkhead box protein P2) glutamatergic projection neurons in the temporal neocortex. Following histological, electrophysiological and transcriptomic characterizations of CellREADR-labeled neurons, we used PatchSeq techniques to cross-validate targeting specificity and perform molecular alignment of the experimentally accessed cells onto published transcriptomic reference atlases. We further performed optical stimulation and calcium imaging in *CALB2* and *FOXP2* populations to probe human cortical microcircuits. Together, our experiments highlight CellREADR as a powerful approach for cell type-specific access, monitoring, and manipulation in human brain circuits, *ex vivo*.

## RESULTS

### Deploying CellREADR to target RNA-defined cell types in ex vivo human cortex

We used an organotypic slice preparation to study CellREADR applications in human cortical tissues (Figure 1A). Live tissue specimens were donated by epilepsy surgical patients and collected from non-diseased areas adjacent to known seizure foci (proximal, non-diseased tissues are minimally removed during surgery to access epileptogenic areas). Donor demographics and specimen details are presented in Table 1.

**Figure 1.**
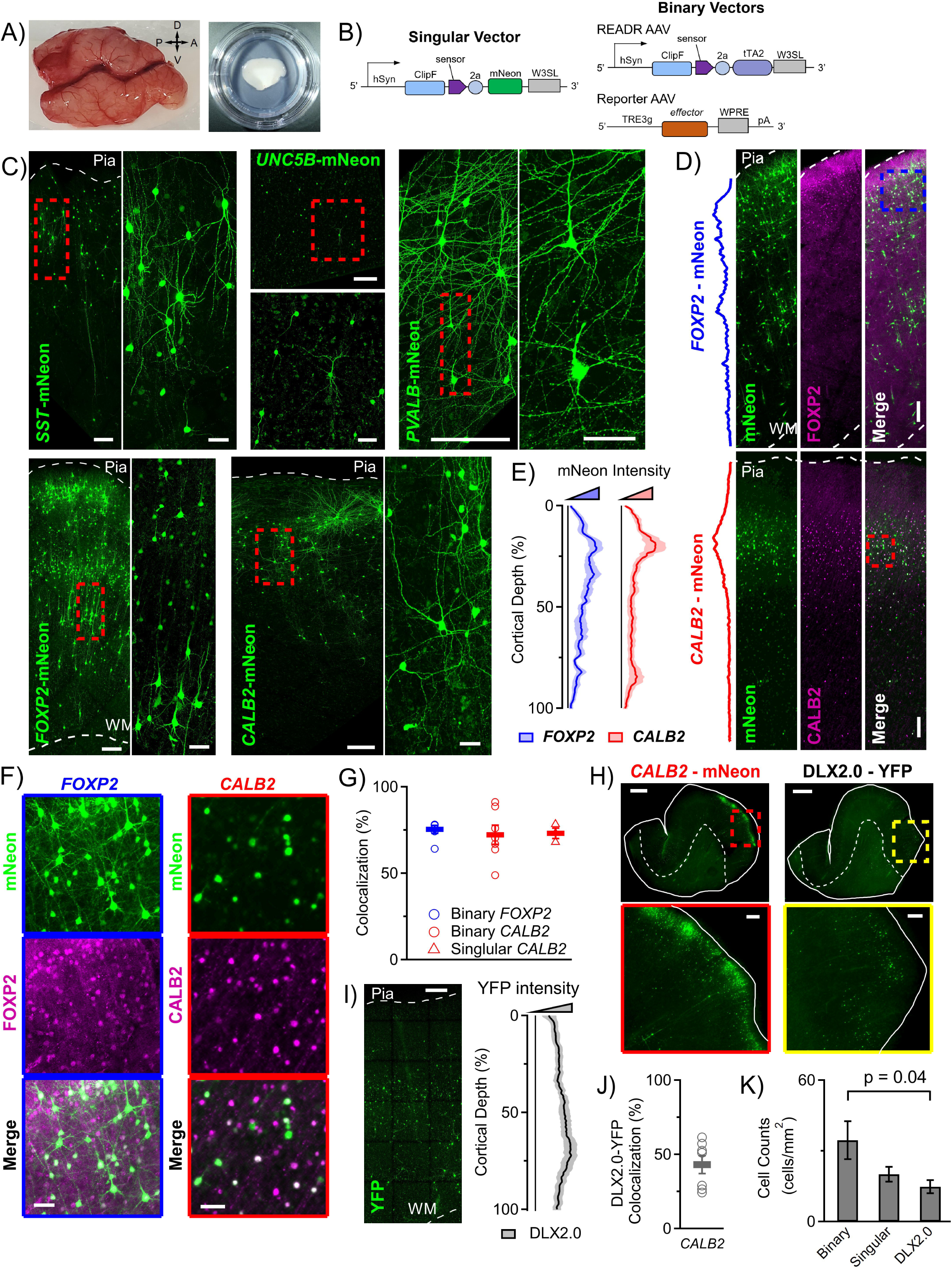
Deploying CellREADR to target RNA-defined cell types in ex vivo human cortex. (A) Human organotypic slice model. Human neocortical tissue specimens (here, middle and inferior temporal gyri; left) were sectioned into 350 µm slices and cultured on semi-permeable membranes for up to 16 days *in vitro* (DIV; right). CellREADR AAV vectors were directly applied to slices on DIV 0. Orientation: A, anterior; D, dorsal; P, posterior; V, ventral. (B) Schematic of CellREADR design. In the single virus design (left), binding of the sensor RNA sequence to the target leads to ADAR-mediated editing of a stop codon positioned between the sensor and the effector, leading to expression of the downstream effector (here, mNeon). The binary system (right) uses a CellREADR virus (top right) to drive expression of tTA2 (Tet-Advanced transactivator) in the target cell. A reporter virus, designed to conditionally express a selected effector molecule (e.g. mNeon, ChIEF, GCamp7f, depending on experiment) under control of the Tet-Responsive Element (TRE), is co-applied. (C) CellREADR targeting of diverse neuronal populations. Binary CellREADRs (mNeonGreen reporter) designed against Somatostatin (*SST*), Unc-5 netrin receptor B (*UNC5B),* Parvalbumin (*PVALB*), Forkhead box protein P2 (*FOXP2)*, and Calretinin (*CALB2*) were applied to slices of human neocortex from at least 3 donors. Slices were immunostained for mNeon at 7 DIV. The CellREADR binary system led to robust expression of mNeon in differing laminar distributions and in cells of various morphologies. Dashed red boxes highlight the area depicted in the accompanying magnified image; pia and white matter (WM) are illustrated with dotted lines. Scale bars: 200 µm in large image and 50 µm in magnified image (D) Histological analysis of binary *FOXP2* and *CALB2* CellREADR targeting. CellREADR mNeon distribution (green, left) and protein expression of target (purple, middle) are depicted at DIV 7. The fluorescence intensity profile to the far left of each image set reveals the distribution of cells targeted by the CellREADR. Dashed red boxes indicate areas depicted in (F); pia and white matter (WM) are illustrated with dotted lines. Scale bars: 200 µm (E) Localization of CellREADR-targeted cells. CellREADR mNeon fluorescence intensity profiles measured across the depth of the cortex (0% depth = pial surface, 100% depth = white matter boundary) averaged from 6 slices (tissues from 3 donors). The solid line shows the mean and the shaded area shows the SEM. *FOXP2*-mNeon was relatively uniform throughout cortical layers, while *CALB2*-mNeon was more restricted to outer layers. (F) Immunohistochemical characterization of CellREADR specificity. Representative images showing colocalization of CellREADR-mNeon with the corresponding target. Scale bars: 50um (G) Quantification of CellREADR specificities, as measured by immunostaining. Each point denotes the specificity of labeling measured from an individual donor’s tissue (*FOXP2* n=5 donors; Binary *CALB2* n=7, Singular *CALB2* n=3). Horizontal bars indicate mean specificity values for each CellREADR. Throughout figures, data are presented as mean ± SEM. (H) Benchmarking of CellREADR efficiency with a human interneuron enhancer virus DLX2.0-YFP. *CALB2* CellREADR and DLX2.0-YFP viruses (rAAV2-retro) were applied to slices cut sequentially from the same neocortical tissue specimen. Tissue was fixed at 7 DIV and immunostained against mNeon or YFP. Slice boundaries are indicated by the solid white line, and the white matter is demarcated by a dashed white line; boxes mark the inset images below. Scale bars: upper images 1 mm, lower images 200 µm (I) Localization of DLX2.0-YFP expression. Representative image illustrating the efficiency and distribution of cellular labeling achieved with a DLX2.0-YFP virus (left), with fluorescence intensity profile (right; averages are from tissues collected from 3 donors) illustrating distribution of cells labeled by the virus. As compared to *CALB2* CellREADR labeling (Figures 1D and 1H), DLX2.0-YFP+ cells were less concentrated in the outer cortex, as would be expected for pan-interneuron targeting by DLX2.0. Scale bar: 250 µm (J) DLX2.0-YFP targeting of the *CALB2* population. Quantification of colocalization of DLX2.0-YFP expression and CALB2, as measured by immunostaining (images not shown); note inter-subject variability in targeting of CALB2 cells (n=7 donors). (K) Efficiency of binary and singular *CALB2* CellREADRs compared to DLX2.0-YFP enhancer virus. Quantification of cells labeled at DIV 7, per unit area (Binary *CALB2*, n=6 donors, Singular *CALB2*, n=3, DLX2.0-YFP, n=3). Note, although the CellREADRs were designed to target only *CALB2* interneurons, rather than multiple subclasses interneurons (as expected for DLX2.0), they labeled more cells (1-way ANOVA with Tukey’s post hoc).

**Table 1.** Donor demographics and specimen utilization.

| Donor ID | Age | Sex | Region of neocortex | Data collected |
| --- | --- | --- | --- | --- |
| TE22 | 8 | M | R parietal | *FOXP2* physiology |
| TE23 | 60 | M | L temporal | *FOXP2* physiology |
| TE29 | 43 | M | L temporal | *CALB2* physiology & morphology |
| TE30 | 52 | M | R temporal | *CALB2* morphology |
| TE31 | 28 | M | R temporal | *CALB2* morphology |
| TE32 | 12 | F | L temporal | *SST* expression |
| TE33 | 45 | M | L temporal | *UNC5B* expression, *CALB2* physiology & morphology |
| TE34 | 19 | F | R temporal | *CALB2* physiology & morphology |
| TE35 | 30 | M | L temporal | *PVALB* expression |
| TE36 | 35 | F | R temporal | *CALB2* expression |
| TE37 | 23 | M | R temporal | *CALB2* physiology |
| TE41 | 42 | M | R temporal | *CALB2* physiology |
| TE44 | 14 | M | L temporal | *CALB2* physiology |
| TE45 | 41 | M | L temporal | *CALB2* physiology |
| TE46 | 43 | F | L temporal | *CALB2* physiology |
| TE47 | 28 | F | L temporal | *FOXP2* expression & physiology, Ca imaging |
| TE48 | 40 | F | R temporal | *FOXP2* physiology |
| TE50 | 35 | F | R temporal | *FOXP2* physiology, Ca imaging |
| TE51 | 29 | F | R temporal | *FOXP2* physiology |
| TE52 | 19 | M | L temporal | *FOXP2* physiology, Ca imaging |
| TE53 | 33 | M | R temporal | *FOXP2* physiology, Ca imaging |
| TE57 | 34 | F | L temporal | Ca imaging |
| TE65 | 47 | F | L temporal | *CALB2* physiology |
| TE66 | 50 | F | R temporal | *FOXP2* physiology |

We designed both singular and binary CellREADR vectors for use in human tissues (Figure 1B). The singular vector system drives expression of the sensor and, upon sense-editing by endogenous ADAR, expression of the effector RNA; this vector was used to directly couple RNA sensing to the translation of an mNeonGreen (mNeon) effector. The binary system was designed to utilize two vectors, one, a READR vector that expressed the transcription activator, tTA2, upon sense-editing, and the other, a reporter vector containing a TRE-driven effector. The binary system enables amplification of effector payload expression and also allows pairing of READR and reporter vectors for differential expression of effectors in separate target populations. The singular vector was used in some histological studies (Figure 1) and a subset of the patch clamp studies (Figures 2 and 3; *CALB2* only), while binary vectors were used for all other experiments, including histological studies (Figure 1; effector, mNeon), patch clamp characterizations of cellular properties (Figures 2 and 3; mNeon), optogenetic interrogations (Figure 4; ChIEF), and calcium imaging (Figure 5; GCaMP7f).

**Figure 2:**
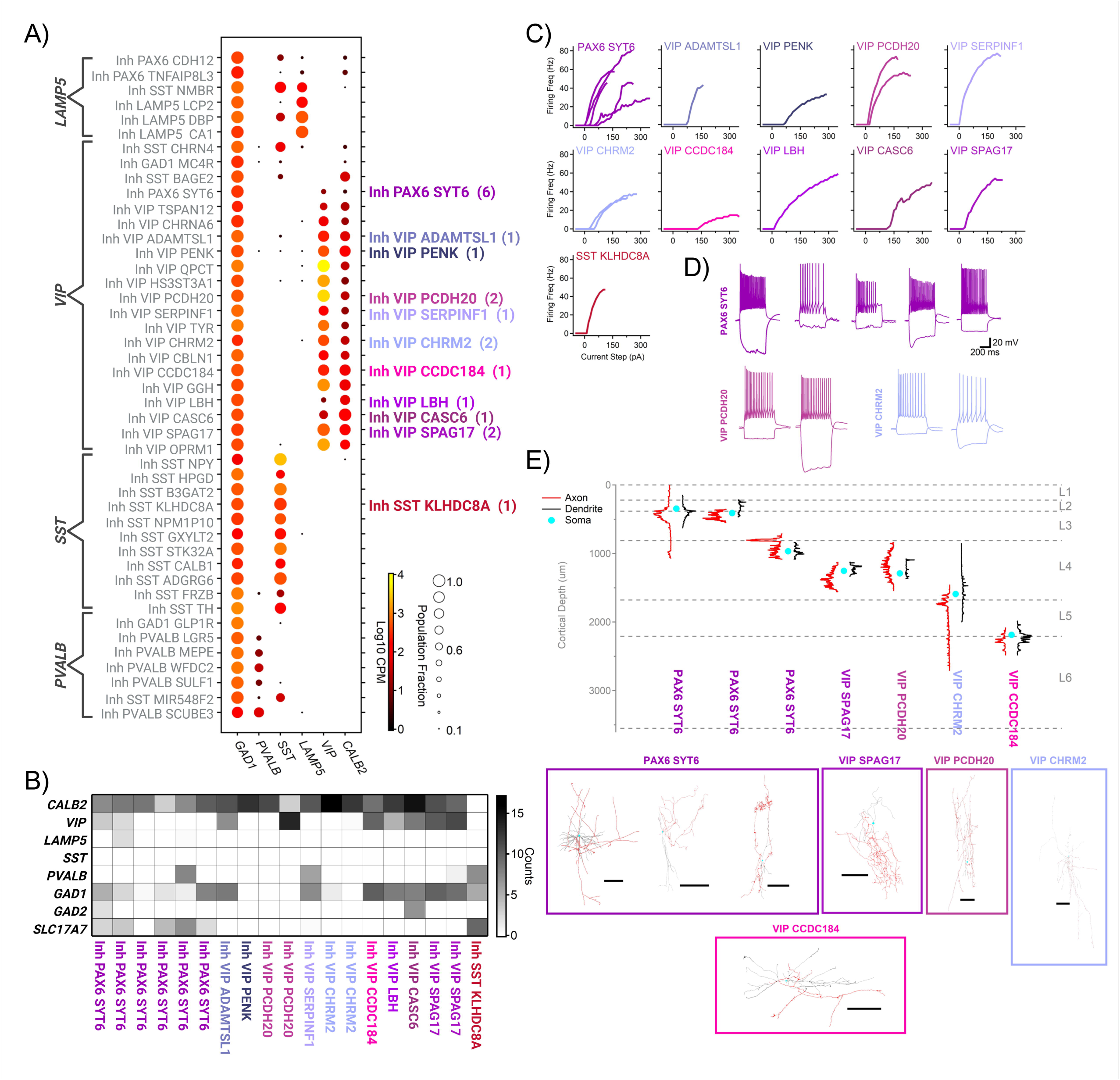
PatchSeq validation of CALB2 CellREADR targeting. (A) Reference gene expression profiles of human MTG interneurons described in a prior study. *CALB2* expression is largely restricted to *VIP* subclass interneurons, indicating that *CALB2* READR is expected to primarily target *VIP* interneurons. Of the 19 *CALB2* READR-targeted cells studied by PatchSeq here, all but 1 cell mapped transcriptomically to the *VIP* subclass (the transcriptomic types of the 19 cells are listed at right; number indicates count of cells mapped to that type) according to the classification scheme adapted from Tasic et al.^77^ and Hodge et al.^7^. (B) Expression of selected genes in *CALB2* READR cells. *CALB2* transcripts were detected in all but 1 cell, which was mapped onto the *SST* subclass (*SST KLDHC8A*). READR-targeted cells rarely expressed canonical markers of interneuron subclasses other than *VIP* (e.g., *LAMP5, SST, PVALB*). Interestingly, 5 of the 6 cells that mapped to the *PAX6 SYT6* type expressed the *SLC17A*7 (vesicular glutamate transporter 1), which is expressed widely across human MTG glutamatergic populations (Supplemental Figure 2A); these cells also expressed the GABAergic marker, *GAD1*. (C) PatchSeq electrophysiological measures of *CALB2* READR-targeted neurons. Input-output curves illustrate the variation in rheobase, gain, and maximum firing frequency across the sampled population. The interrelationship between gene expression and physiological properties is further illustrated in Supplemental Figure 3. (D) Variation in electrophysiological properties across and within interneuron transcriptomic types. Membrane responses to depolarizing and hyperpolarizing currents from cells that were mapped to the same transcriptomic types. Cells that mapped to the *PAX6 SYT6* type exhibited variation in the frequency and accommodation of action potential firing and also exhibited varying responses to hyperpolarization. Scale bars: 20 mV, 200 ms (E) Morphologies of *CALB2* READR-targeted cells from the PatchSeq subset. The positions (within cortical mantle) of reconstructed neurons are illustrated at top. Blue circles indicate locations of the cell somata (not to scale), black profiles indicate the vertical and horizontal densities of cell dendrites, and red profiles indicate the vertical and horizontal densities of cell axons. Cortical depth is indicated on the left vertical axis; average cortical layer boundaries are shown on the right vertical axis. Complete cell tracings are shown at bottom. Supplemental Figure 5 shows the morphologies of additional *CALB2*- and *FOXP2*-targeted cells. Scale bars: 200 µm

**Figure 3:**
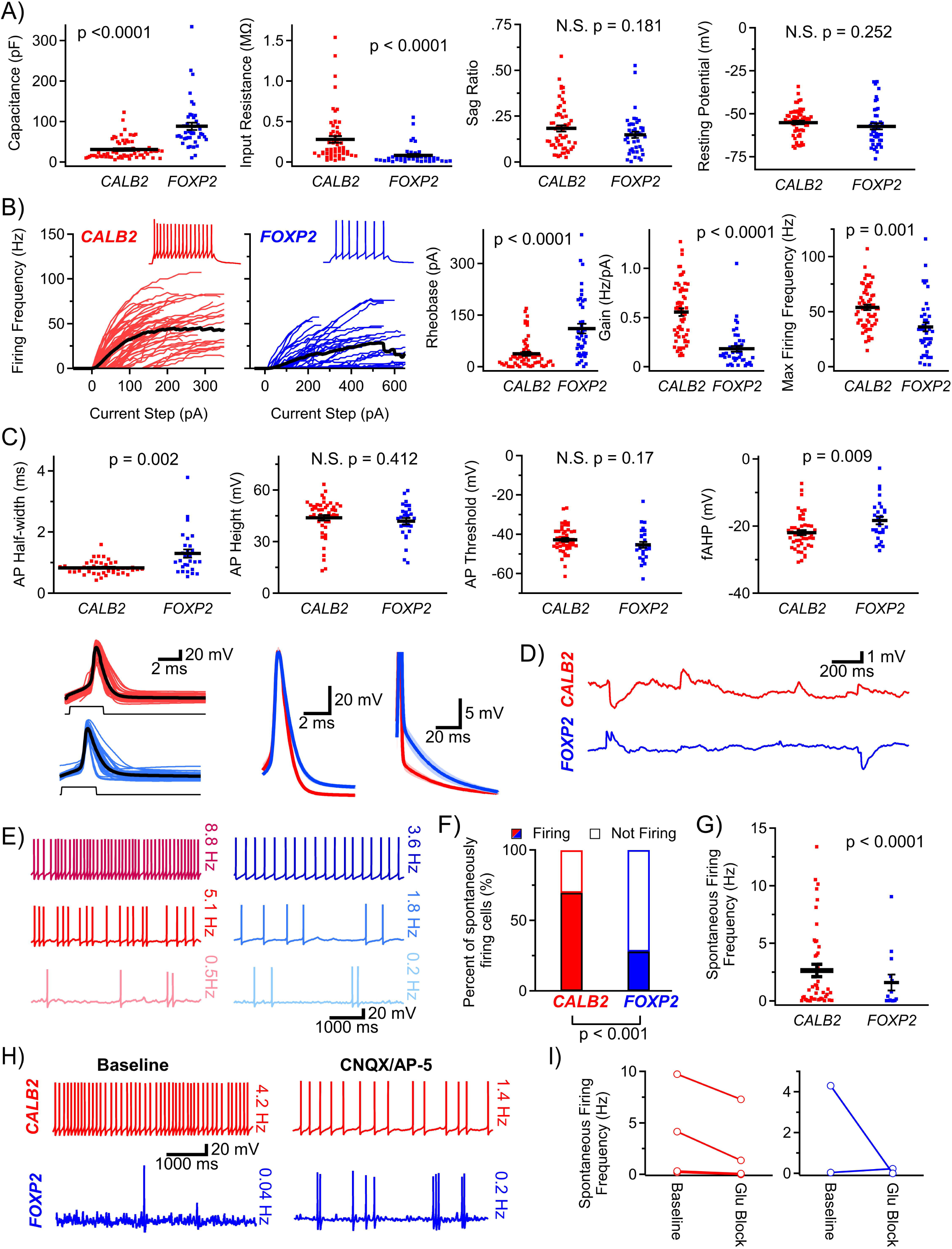
CALB2- and FOXP2-targeted neurons exhibit distinct physiological profiles and spontaneous activities. (A) Passive membrane properties of cells targeted by *CALB2* (n=60 cells) and *FOXP2* CellREADRs (n=43). Membrane capacitance and input resistance, but not hyperpolarizing sag ratio or resting membrane potential, differed statistically between the two target populations. Black bars indicate mean and SEM (Mann-Whitney U-test for non-normal distributions, t-test for normal distributions). (B) Active properties of cells targeted by *CALB2* (n=60 cells) and *FOXP2* CellREADRs (n=43). Firing responses of cells in response to current injections, with frequency plotted against current step. Insets show action potentials elicited by current injections at 2X rheobase; these examples were selected to represent the mean input-output curves for each cell type (black). The rheobase (current that initiated firing), gain (initial slope), and max firing frequency all differed statistically between *CALB2* and *FOXP2*-targeted populations (Mann-Whitney U-test for non-normal distributions, t-test for normal distributions). (C) Action potential (AP) properties of cells targeted by *CALB2* (n=60 cells) and *FOXP2* CellREADRs (n=43 cells). Action potentials were elicited with a 2ms current step. Action potential properties from different stimulation regimes are presented in Supplemental Figure 4. Action potential half-width was measured at 50% of peak voltage (height) and was found to be smaller in *CALB2*-targeted cells. Action potential height and threshold did not vary between target cell groups. The fast afterhyperpolarization (fAHP) following the AP was larger in *CALB2*-targeted cells. Action potentials were aligned on their rising phase to compare AP waveforms between target cell groups (lower panel). Group means are illustrated by the black lines. *CALB2*-targeted cells exhibited a shorter AP duration and a smaller afterdepolarization than *FOXP2*-targeted cells (Mann-Whitney U-test for non-normal distributions, t-test for normal distributions). Scales bars: left 20 mV, 2 ms; center 20 mV, 2 ms; right 5 mV, 20 ms (D) Subthreshold synaptic activity and suprathreshold firing were observed in targeted cells (held at resting membrane potential with 0 pA current). Excitatory and inhibitory postsynaptic potentials (EPSPs and IPSPs, respectively) were detected in both *CALB2-* and *FOXP2-*targeted populations. Representative traces demonstrating EPSPs and IPSPs in cells of both populations are shown. Scale bars: 1 mV, 200 ms (E) Recorded cells exhibited spontaneous firing from their resting potentials. Examples of spontaneous firing activity observed in *CALB2*- (red, n=3 cells) and *FOXP2*-targeted cells (blue, n=3) are illustrated, with average firing frequencies (observed over a 3-minute period) indicated to the right of the traces. Scale bars: 20 mV, 1000 ms (F) *CALB2*-targeted cells (n= 66 cells) were more likely than *FOXP2*-targeted cells (n=50) to exhibit spontaneous firing in the slice preparation (Chi-squared test). (G) Spontaneous activity, measured as the frequency of spontaneous action potential firing, differed between *CALB2- and FOXP2*-targeted populations (Mann-Whitney U test). (H) Traces demonstrating spontaneous action potential firing from *CALB2*- (red) and *FOXP2*-targeted cells (blue) in either baseline conditions (left) or in the presence of glutamatergic blockers (right). Scale bars: 20 mV, 1000 ms (I) Glutamatergic blockers (CNQX/AP-5) reduced spontaneous firing frequency in most cells (*CALB2*, n=6 cells; *FOXP2*, n=2), but did not eliminate firing in every cell.

**Figure 4:**
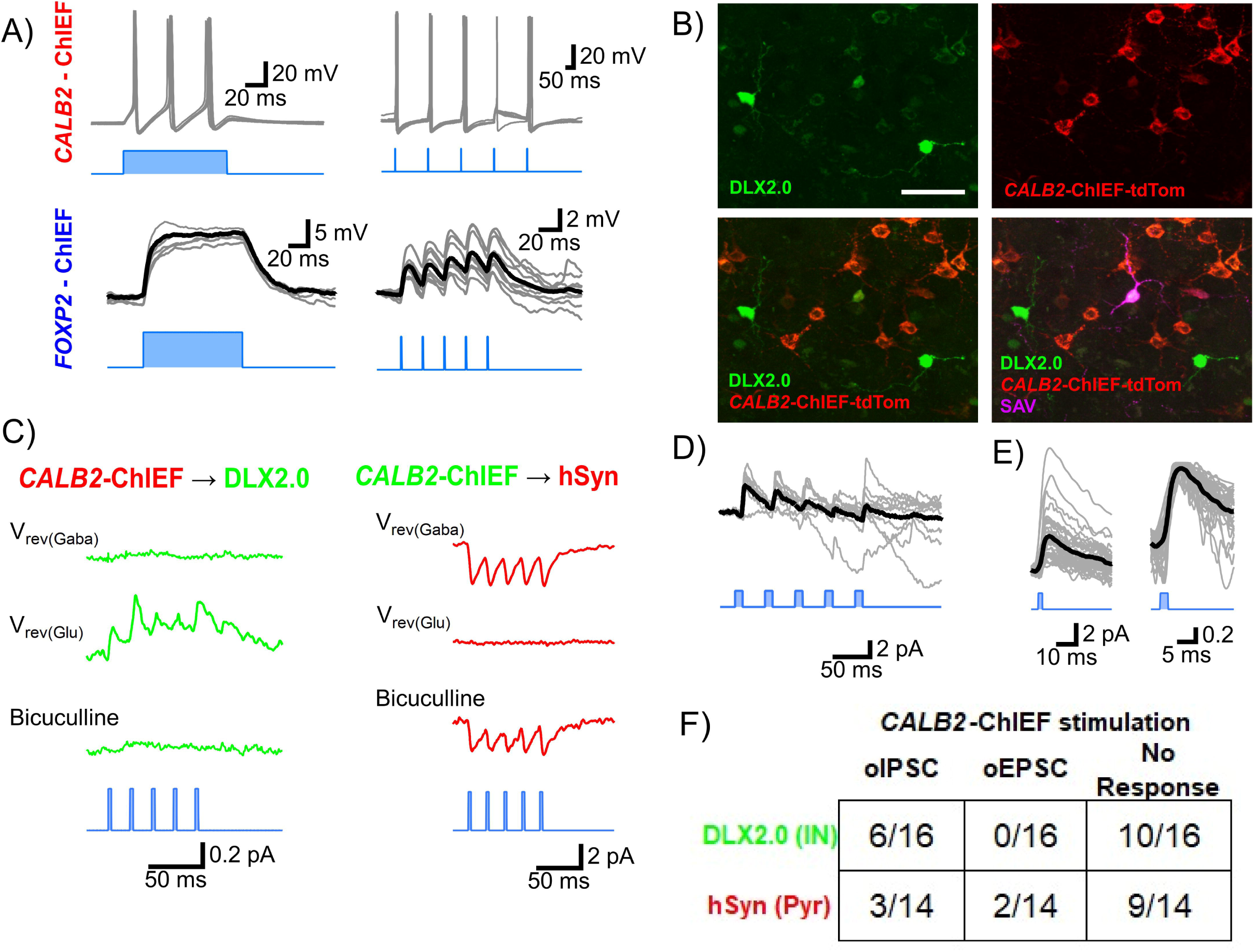
Cell type-specific optical manipulation of human neurons with CellREADR. (A) CellREADR expression of an optogenetic effector, ChIEF, enables optogenetic control of action potential firing in *CALB2*-but not *FOXP2*-targeted cells. *CALB2-*ChIEF neurons fired action potentials (APs) in response to 100 ms, 642 nm light pulses (left) or trains of 2 ms pulses (6/6 cells fired action potentials in response to both stimuli, with some failures at the 4th stimulus). Light stimuli depolarized *FOXP2*-ChIEF, but did not drive AP firing (0/10 cells fired action potentials in response to stimuli; grey traces, individual trials from a single cells, black traces, averages). Scale bars: 20mV, 20ms, 50ms (B) Representative targeting scheme for optical interrogation of *CALB2* synaptic connectivity. Putative postsynaptic cells were labeled with a DLX2.0-YFP virus (left top, green). A binary CellREADR vector driving *CALB2*-ChIEF-tdTomato was used to activate presynaptic cells (right top, red). During optical stimulation of the ChIEF-tdTomato^+^ population, patch clamp recordings were made from YFP^+^/ChIEF-tdTomato^-^ cells; at the end of recordings, biocytin-filled patched cells were recovered (right bottom, purple, SAV). Scale bar: 50 µm (C) Postsynaptic currents elicited by CALB2-ChIEF activation. Left, voltage clamp recordings (average of 10 trials in all cases) made from a DLX2.0-YFP^+^/ChIEF-tdTomato^-^ neuron during optical stimulation of the CALB2-ChIEF population. Stimulation elicited optical postsynaptic currents (oPSCs), which were present at the glutamate reversal potential (V_rev(Glu)_) but absent at the GABA reversal potential (V_rev(GABA)_), indicating that they were mediated by synaptically released GABA. Bicuculline was applied to verify observed oIPSCs were GABAergic. In DLX2.0-YFP^+^/ChIEF-tdTomato^-^ cells, oPSCs were never observed at V_rev(GABA)_. Right, averaged traces recorded from a hSyn-mCherry^+^ pyramidal neuron during optical stimulation of the *CALB2*-ChIEF population. In the recording shown here, oPSCs were detected at V_rev(GABA)_, but not V_rev(Glu)_, indicating the cell received glutamatergic inputs from *CALB2*-ChIEF cells. These inputs were not blocked by bicuculline. No cell exhibited oPSCs at both V_rev(GABA)_ and V_rev(Glu)_. Scale bar: left 0.2 pA, 50 ms; 2 pA, 50ms (D) Failure rate of optically triggered synaptic currents. To illustrate trial-to-trial variability in oPSC responses, 10 trials from a DLX2.0-YFP^+^/ChIEF-tdTomato^-^ cell are shown at V_rev(Glu)._ The first light pulse had a failure rate of 10% (1/10), while the fourth light pulse had a failure rate of 40% (4/10; averaged response, black). Scale bar: 2 pA, 50 ms (E) Waveforms of optically-evoked postsynaptic responses. Optical PSCs measured at V_rev(Glu)_ from DLX2.0-YFP^+^/ChIEF-tdTomato^-^ interneurons and hSyn-mCherry+ pyramidal neurons were aligned to the light pulse and overlaid to illustrate the variability in amplitudes (left). All oPSCs were normalized to the peak current (right). The monophasic rise time and small jitter relative to the 2 ms light stimulus indicate that PSCs were monosynaptic events directly triggered by the optical pulse (averaged response, black). Scale bar: left 2 pA, 10 ms; 0.2, 5 ms. (F) Quantification of postsynaptic responses to *CALB2*-ChIEF activation. Recordings were made from DLX2.0-YFP^+^/*CALB2*-ChIEF^-^ interneurons during *CALB2*-ChIEF activation. Optical inhibitory PSCs (oIPSCs, oPSCs detected at V_rev(Glu)_) were observed in 6/16 cells (37.5%), and optical excitatory PSCs (oEPSCs, oPSCs detected at V_rev(GABA)_) were not detected. Recordings were also made from hSyn-tdTomato^+^ pyramidal neurons during *CALB2*-ChIEF activation. Optical IPSCs were detected in 3/14 cells (21.4%), and oEPSCs were detected in 2/14 cells (14.3%).

**Figure 5:**
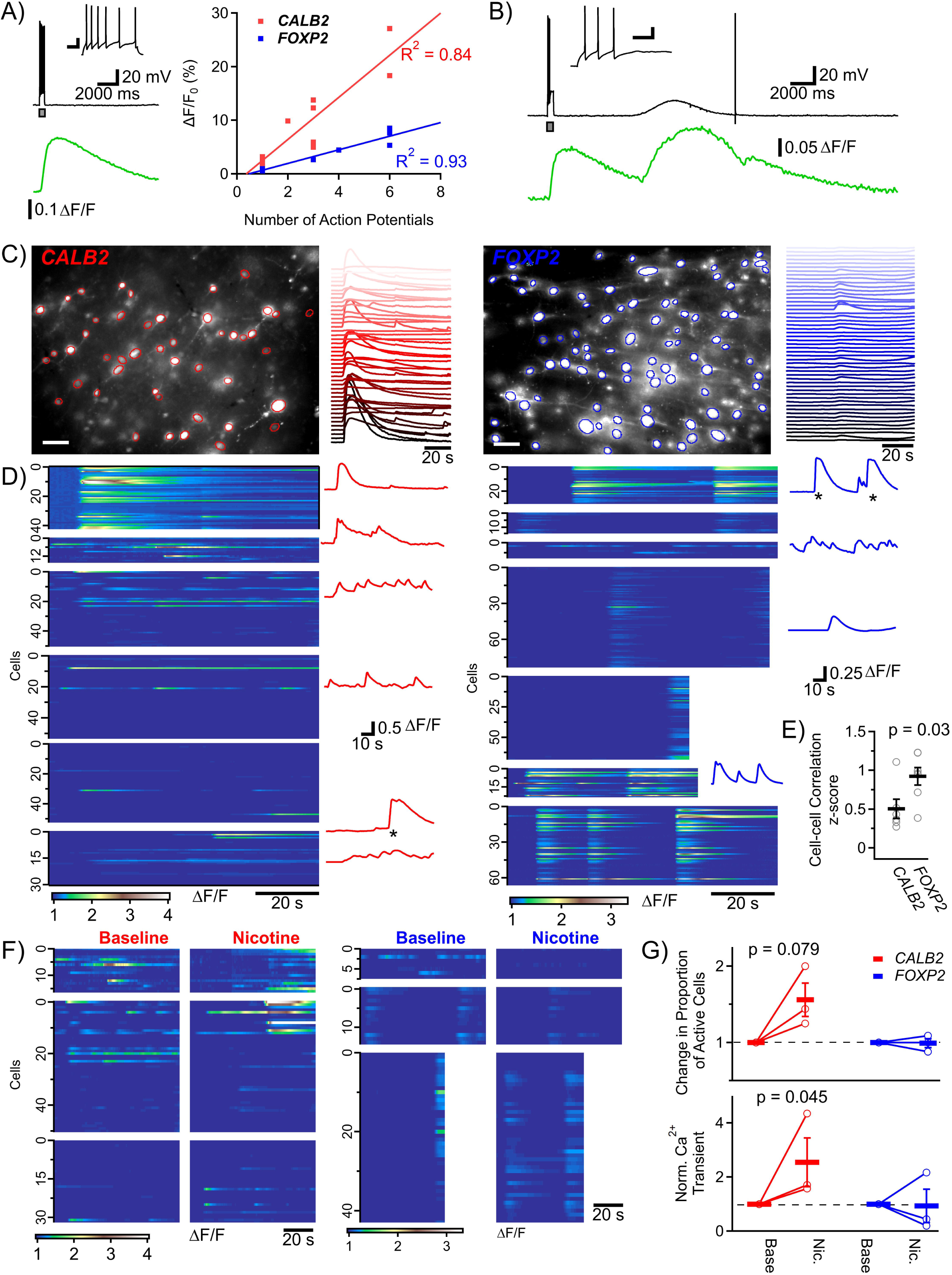
Cell type-specific monitoring of population activity with CellREADR. (A) Correlation between action potential firing and GCamp7f signals. Action potentials (black) were elicited by current injections in single cells expressing either *CALB2*-GCamp7f (n=4 cells) or *FOXP2*-GCamp7f (n=4; top left, current injection is represented by gray box; the upper inset shows the voltage trace at an expanded time scale to show the individual action potentials (APs)). Calcium transients (green) were concurrently imaged at the cell soma and proximal dendritic compartment. Traces show 6 APs and resultant GCaMP ΔF/F signal measured from a *CALB2*-GCamp7f cell. The relationship between the baseline-subtracted change in fluorescence and AP firing (right) was established by varying the current injection, counting APs, and fitting the ΔF/F to action potential counts with a linear function (*CALB2*: R^2^ = 0.84; *FOXP2*: R^2^ = 0.93). Scale bar: inset 20 mV, 100 ms; 20 mV, 2000 ms (B) Calcium imaging detects subthreshold events as well as action potentials. Recording of a *CALB2*-GCaMP7f demonstrating three evoked action potentials (black trace, at grey box) followed by a spontaneous subthreshold depolarization then a single spontaneous action potential. The calcium transient for these three events (green) is depicted; the rise times for each event were as follows: Tau_3AP_ = 390 ms, Tau_Depolarization_ = 1760 ms, Tau_1AP_ = 202 ms. Scale bars: inset 20 mV, 100 ms; 20 mV, 2000 ms (C) Maximum projection of *CALB2*-GCamp7f (left, red) and *FOXP2*-GCamp7f (right, blue) neurons measured in baseline conditions (standard artificial cerebrospinal fluid). Background-subtracted fluorescent profiles of cell somata were collected from 90 s video stacks (red and blue traces). CellREADR typically labeled 10-50 cells within a field of view. Scale bars: 50 µm, 20 s (D) Color maps depicting population calcium signal dynamics. Calcium signals from slices labeled with *CALB2*-GCamp7f (left, red; n=6 slices) or *FOXP2*-GCamp7f cells (right, blue n=7) are represented by a color map (intensity scale at bottom; each map represents a recording performed on a single slice, slices were collected from 4 donors). Spontaneous activity levels varied between slices. Calcium signal traces from selected individual cells are depicted to the right of the color maps; most transients were small-amplitude, fast-rising events, similar to the transients produced by APs in (A). Large-amplitude, slow-rising transients (marked with asterisks) similar to the subthreshold events in (B), were also observed. Scale bars: population maps 20 s, individual traces 10 s (E) Correlated calcium events in slice populations. The correlation of cellular *CALB2*-GCaMP7f (n=6 slices) or *FOXP2*-GCaMP7f (n=7) calcium signals was measured within each slice and then compared between groups (*CALB2-* and *FOXP2*-targeted populations). Calcium signals were more strongly correlated in the *FOXP2*-targeted population (Unpaired t-test). (F) Calcium signal responses to nicotinic acetylcholine receptor activation. Nicotine (300 μM) was applied to a subset of slices following imaging of baseline activity. Each panel represents a ROI in a single slice labeled with either *CALB2*-GCaMP7f (left, red; n=3 slices) and *FOXP2*-GCaMP7f (right, blue; n=3). Scale bar: 20 s (G) Nicotine application did not change the proportion of active cells within the labeled populations (*CALB2*, p=0.08; *FOXP2*, p=0.999 by Fisher exact test). However, nicotine application increased the area of measured calcium events, normalized to baseline areas in *CALB2*-targeted cells (p = 0.045 by 2-way ANOVA with Fisher’s LSD post-hoc). Dashed line on both graphs represents unity.

We first produced CellREADR sensors for various interneuron and projection neuron marker genes (*VGAT* (n=2 sensors), *LAMP5* (n=1), *CALB2* (n=3)*, SST* (n=2), *PVALB* (n=3)*, UNC5B* (n=1)*, CUX1* (n=1)*, FOXP2* (n=2); a total of 15 sensors) and used them in binary vector designs to express mNeon in cells of the human cortex. In screening these designs on human tissues, we observed READR activity with 13 sensors. Labeled cells exhibited diverse morphological features that were consistent with glutamatergic and GABAergic neurons, depending on the sensor (example labeling, Figure 1C). We elected to pursue more detailed investigations using two screened sensors, one for *FOXP2* and the other for *CALB2*, given their robust expression in human slices and prior data suggesting these RNAs may be expressed in cell populations that confer or exhibit human specializations.

*FOXP2* is implicated in human language disorders^51,52^, aberrant song learning in songbirds^53^, and impairments in fine motor learning in mice^54^. In rodents, *FOXP2* expression is restricted to cortical excitatory neurons in deep layers^55^, however, *FOXP2* is expressed more widely across cortical lamina in humans^48,56^. Consistent with this, after applying a binary *FOXP2* CellREADR system to neocortical slices, we observed mNeon epifluorescence in pyramidal cells of both deep and superficial cortical layers (Figures 1C-E). To assess the specificity of *FOXP2* CellREADR targeting, we performed immunohistochemistry to quantify co-labeling of READR mNeon with cellular FOXP2 7 days after virus application (Figure 1F). Across tissues from 5 subjects, the mean specificity of the *FOXP2* READR labeling (percentage of mNeon+ cells co-labeled with FOXP2) was 75.4 ± 2.2% (Figure 1F-G).

In the human cortex, *CALB2* expression is largely restricted to supragranular GABAergic interneurons that originate from the caudal ganglionic eminence (CGE), primarily *VIP* subclass interneurons^57–59^. There is evidence for an increased abundance of CGE interneurons (versus medial ganglionic eminence interneurons) in humans and non-human primates, as compared to mice^3,7,9^, with some studies suggesting a specific expansion of *CALB2* populations^57,60,61^. Using a binary *CALB2* READR system in human neocortical slices, we observed a pattern of mNeon labeling concordant with prior descriptions of *CALB2* expression. Compared to *FOXP2* READR labeling, *CALB2* READR expression was generally observed in superficial cortical layers (Figures 1C-E). *CALB2* READR-labeled cells also exhibited morphologies distinct from *FOXP2* READR-labeled cells, with small somata and vertically oriented, bipolar or multipolar dendritic processes (Figure 1C). The specificity of the binary *CALB2* READR transduction, as tested by immunohistochemistry in tissues from 7 subjects, was 72.2 ± 5.5%; specificity of the singular *CALB2* READR was 73.1 ± 3.0% (3 subjects; Figures 1F-G). Quantification of immunostaining signals in FOXP2 and CALB2 cells indicated that CellREADR sense-editing did not disturb CALB2 expression in targeted cells, whereas FOXP2 expression was found to be slightly lower in cells targeted by the *FOXP2* READR (Supplemental Figure 1).

To benchmark targeting by the *CALB2* CellREADRs, we compared READR labeling to that achieved by an enhancer virus that has been recently used for human pan-interneuron targeting, DLX2.0^20^ (Figure 1H). Consistent with prior evidence that DLX2.0 transduces broadly across GABAergic cell subclasses^33^, 43.1 ± 6.0% of DLX2.0-YFP+ cells co-expressed CALB2 (Figure 1J),. Compared to *CALB2* CellREADR labeling, which was noted primarily in supragranular layers, DLX2.0 virus expression was also more evenly distributed across the depth of the cortical gray matter (Figure 1I). Although the binary and singular *CALB2* CellREADR vectors theoretically targeted a more restricted population of interneurons, we found that they labeled greater numbers of cells than the DLX2.0 virus, suggesting their greater targeting efficiency (Figure 1K; Binary *CALB2*-READR, 34.5 ± 8.0 cells/mm^2^, Singular *CALB2*-READR, 20.1 ± 3.5 cells/mm^2^, DLX2.0-YFP, 14.8 ± 2.8 cells/mm^2^; 1-way ANOVA, p = 0.04).

### PatchSeq validation of CALB2 CellREADR targeting

To further validate CellREADR targeting and gain a multimodal characterization of some of the cells accessed by the *CALB2* READR, we performed READR-targeted PatchSeq on a limited set of cells. Four to 12 days after the application of singular or binary *CALB2* READR vectors, we targeted mNeon+ cells for patch clamp electrophysiology, biocytin filling, and nuclear extraction. After performing RNA-seq on the nuclear aspirate, we applied an open sequencing analysis pipeline^62^ to map queried cells onto a reference transcriptomics-based taxonomy of human middle temporal gyrus (MTG) neurons^59^ (Figure 2A, Supplemental Figure 2A).

As expected, our RNA-seq analysis detected *CALB2* transcripts in nearly all mNeon+ patch-targeted cells (18/19, 94.7%; Figure 2B). Notably, this exceeded the specificity of *CALB2* READR targeting measured by immunohistochemistry (Figure 1G), suggesting that incomplete sensitivity of CALB2 immunostaining may have underestimated the specificity of the *CALB2* READR. These RNA-seq results also indicate the CellREADR sense-edit mechanism did not grossly disrupt expression of the *CALB2* target gene. Based on existing transcriptomics data^59^ (Figure 2A), which demonstrate that *CALB2* is expressed almost exclusively by *VIP* subclass interneurons, we hypothesized that *CALB2* READR-targeted cells would also express the interneuron class marker, *GAD1*, as well as *VIP*, but not markers of other cortical interneuron subclasses, such as *LAMP5*, *SST*, and *PVALB*. In accordance, transcripts of *GAD1* were detected in the majority of sampled cells (14/19, 73.7%), while *VIP* was detected in a somewhat smaller fraction (9/19; 47.4%); by contrast, transcripts of *LAMP5, SST, PVALB* were rarely observed in *CALB2* READR-targeted cells (Figure 2B). Looking beyond single gene expression, essentially all cells (18/19; 94.7%) mapped transcriptomically to types within the *VIP* subclass of MTG interneurons (Figures 2A and 2B). Of cells that mapped to the *VIP* subclass, one-third (6/18; 33.3%) were categorized as *PAX6 SYT6* type, while the remaining two-thirds (12/18; 66.7%) mapped to other *VIP* types. In total, 10/21 previously annotated transcriptomic types of *VIP* interneurons^63^ were represented amongst the 19 cells sampled by *CALB2* READR targeting. These results together indicate that CellREADR specifically targeted *CALB2* populations and, thereby, effectively accessed *VIP* interneurons (Supplemental Figure 2B). Expected transcriptomic alignment of *CALB2* READR-targeted cells onto this existing database (which was produced without CellREADR) suggest CellREADR did not significantly alter target cell transcriptomes, consistent with immunohistochemistry data provided in Supplemental Figure 1 and results we have presented in a previous study^48^.

Electrophysiology data from 17/19 cells studied with RNAseq passed quality checks, and the morphologies of 7 cells were recovered for tracing. *CALB2* READR-targeted neurons largely showed a steep gain relationship between injected current and firing frequency, with many cells exhibiting firing frequencies exceeding 40Hz (Figure 2C). Qualitatively, cells that clustered within the same transcriptomic type nevertheless exhibited differences in their electrophysiological features. For example, within READR-targeted cells identified as *PAX6 SYT6* type, we observed two distinct patterns of input-output responses (Figure 2C), as well as similarly variable responses to hyperpolarization (Figure 2D). These basic phenotypic observations were consistent with recent human interneuron PatchSeq data^33^, and generally align with prior observations that cell transcriptomic types do not necessarily express type-specific physiological features. As expected, however, we did observe some relationships between ion channel subunit expression and membrane physiology^38^. The expression of BK channel subunits, for example, correlated with both the depth of the fast afterhyperpolarization and the action potential duration, as would be expected given the function of this channel^64,65^ (Supplemental Figure 3).

Morphologically, reconstructed neurons were located in Layers 2-5 and exhibited diverse dendritic and axonal distributions. Six of 7 cells projected dendrites vertically, while the other recovered cell’s dendrites were oriented horizontally (Figure 2E). These morphologies are likewise generally consistent with *VIP* subclass morphologies^33^.

### CALB2- and FOXP2-targeted neurons exhibit distinct physiological profiles and spontaneous activities

After gaining evidence for CellREADR specificity through immunostaining and sequencing analyses, we proceeded to a more detailed physiological characterization of neurons targeted by *CALB2* and *FOXP2* READRs. Applying a panel of electrophysiological readouts used to construct prior human cellular atlases^33,43,66^, we recorded electrophysiological properties from 60 *CALB2* READR-targeted cells (n=8 subjects) and 43 *FOXP2*-targeted cells (n=7 subjects). We first measured intrinsic membrane properties and noted group differences in membrane capacitances (*CALB2*: 31.72 ± 3.35 pF; *FOXP2*: 88.46 ± 9.05 pF, Mann-Whitney p<0.0001) and input resistances (*CALB2*: 0.280 ± 0.041 MΩ; *FOXP2*: 0.082 ± 0.018 MΩ, Mann-Whitney p<0.0001; Figure 3A). Hyperpolarizing sag ratios (*CALB2*: 0.18 ± 0.02; *FOXP2*: 0.15 ± 0.02) and resting membrane potentials (*CALB2*: -55.07 ± 1.04 mV; *FOXP2*: -57.30 ± 1.77 mV) did not differ between targeted cells (Figure 3A). We also characterized active membrane properties by quantifying cellular input-output responses (rheobase, gain, and maximal firing frequency; Figure 3B). *CALB2*-targeted cells showed more variability in the shapes of their input-output functions, with many neurons exhibiting low rheobase, high gain, and high maximal firing frequency (41/59 (65%) with maximum firing frequency >40Hz), and a smaller subset demonstrating high rheobase, low gain, and low maximal firing frequency. *FOXP2*-targeted cells were of more uniform firing behaviors, with relatively shallow gain slopes and maximal firing frequencies typically under 40Hz (11/46 (24%) with maximum firing frequency >40Hz). Statistically, rheobases (*CALB2*: 37.9 ± 5.9 pA; *FOXP2*: 110.8 ± 14.0 pA, paired Mann-Whitney p<0.0001), gains (*CALB2*: 0.55 ± 0.04 Hz/pA; *FOXP2*: 0.18 ± 0.03 Hz/pA, paired t-test, p<0.0001), and maximal firing frequencies (*CALB2*: 53.94 ± 2.57Hz, *FOXP2*: 36.15 ± 4.03Hz, Welch’s t-test p=0.001) differed between *CALB2-* and *FOXP2*-targeted cells (Figure 3B).

We performed brief current injections (2 ms) to measure evoked single action potential parameters, including threshold, peak, half-width, and fast afterhyperpolarization from targeted cells (Figure 3C). All single action potentials from the *CALB2* and *FOXP2* cells were aligned on the rising slope and averaged. *CALB2*-targeted cells had narrower action potentials (half-width, *CALB2*: 0.83 ± 0.03 ms; *FOXP2*: 1.30 ± 0.13 ms, Welch’s t-test p=0.002; duration (not shown), *CALB2*: 1.99 ± 0.08 ms; *FOXP2*: 2.78 ± 0.44 ms, Mann-Whitney p=0.003). The *CALB2* population had a larger fast afterhyperpolarization (fAHP; *CALB2*: -21.9 ± 0.75 mV; *FOXP2*: -18.29 ± 1.13 mV, Welch’s t-test p=0.009); a subset of the *FOXP2*-targeted neurons were notable for their pronounced afterdepolarizations. Neither the action potential thresholds (*CALB2*: -42.77 ± 0.98 mV; *FOXP2*: -45.36 ± 1.53 mV) nor the action potential heights (*CALB2*: 43.95 ± 1.60 mV; *FOXP2*: 41.98 ± 1.92 mV) statistically differed between the targeted populations. In an alternate stimulation paradigm (500 ms current injections), we also observed these differences in action potential waveforms between *CALB2*- and *FOXP2*-targeted neurons (Supplemental Figures 4A and 4B). The observed differences in intrinsic properties, firing phenotypes, and evoked action potential dynamics also indicate the *CALB2* and *FOXP2* READRS targeted distinct populations of interneurons and projection neurons, respectively^67,68^. Additionally, and consistent with these findings and the results in Figure 1, reconstructions of the patched cells also demonstrated corresponding morphological differences between the targeted populations (Supplemental Figure 5).

Consistent with previous results^33^, we did not find evidence that intrinsic properties measured at later time points (8-12 days after slice preparation) differed from data collected at earlier points (Supplemental Table 1). Likewise, the *CALB2* READR-targeted cells exhibited physiological properties that resembled those of *VIP* subclass human interneurons collected with an alternate targeting approach^33^ (either without the use of viruses, or with a DLX2.0 virus; Supplemental Table 2). These data indicate that over the period of slice culture CellREADR also did not grossly disturb cellular physiology.

We also used patch clamp to measure activity from *CALB2*- and *FOXP2*-targeted cells at rest, in order to assess synaptic inputs and cellular activity in the slice network. In both target populations, we observed excitatory and inhibitory postsynaptic potentials (EPSPs and IPSPs; Figure 3D), as well as action potential firing (Figure 3E). *CALB2*-targeted neurons often fired from resting membrane potential, whereas a smaller fraction of *FOXP2*-targeted cells exhibited this behavior (*CALB2*: 45/66 cells (69.7%), *FOXP2*: 14/50 cells (28.0%); Fisher’s exact test p<0.001; Figure 3F). Spontaneous activity, measured as the frequency of spontaneous action potential firing, differed between *CALB2- and FOXP2*-targeted populations (*CALB2*: 2.65 ± 0.53 Hz; *FOXP2*: 1.60 ± 0.69 Hz, Mann-Whitney p<0.0001, Figure 3G). The properties of spontaneous action potentials did not differ between groups (Supplemental Figures 4C and 4D). Overall, these observed differences in spontaneous activity were consistent with our intrinsic physiological measures (Figure 3), which indicated the *CALB2*-targeted population is more excitable, with lower rheobase and higher input resistance.

In a subset of spontaneous action potential recordings, we monitored firing before and after application of glutamatergic blockers (CNQX/AP-5; Figure 3H). Glutamatergic blockade resulted in increased (1/6 cells), abolished (3/6), or partially reduced firing (2/6; Figure 3I). As some cells exhibited spontaneous firing while under excitatory blockade, the baseline activity measured at rest reflected a mixture of synaptic recruitment by excitatory inputs as well as spontaneous action potential generation.

### Cell type specific optical manipulation of human neurons with CellREADR

We next explored CellREADR functionality for cellular manipulation and interrogation of human microcircuits. We designed a binary vector system consisting of either *CALB2*-tTA2 or *FOXP2-*tTA2 paired with a TRE-driven channelrhodopsin variant, ChIEF^69^, and then delivered viruses to neocortical slices. We first used patch clamp to assess optically evoked action potential firing in *CALB2*-ChIEF and *FOXP2*-ChIEF cells. In all *CALB2-*targeted cells (n=6), we observed reliable, light-entrained action potentials (Figure 4A); each of these cells was able to follow light pulses with action potentials at frequencies 25-50Hz (data not shown). By contrast, optical stimulation of *FOXP2*-ChIEF cells evoked membrane depolarizations of 2-9mV, but it did not elicit action potentials (n=10 cells; Figure 4A). These differences in light responsiveness were somewhat expected based on our intrinsic physiological characterizations of these populations (Figure 3), which indicated that *FOXP2*-targeted cells are less excitable than *CALB2*-targeted cells. We thus focused on additional optogenetic experiments with *CALB2* READRs.

It has been reported that calretinin interneurons project onto local inhibitory and excitatory populations in rodent cortex^70^ and hippocampus^71^. In non-human primate visual cortex CALB2-immunostained cells have been found to target glutamatergic neurons in deeper layers^72^. Here, to functionally interrogate the synaptic targets of *CALB2* neurons in human neocortex, we recorded from pyramidal neurons and interneurons while optically stimulating *CALB2*-ChIEF cells. Putative postsynaptic cells were labeled with complementary DLX2.0-YFP (Figure 4B) or hSyn-mCherry AAV vectors to label YFP+ interneurons and mCherry+ pyramidal neurons, respectively, for patch clamp recordings (Figure 4B). At short latencies following *CALB2*-ChIEF stimulation, we observed optical inhibitory PSCs (oIPSCs) in slice interneurons and pyramidal neurons, which were absent at the reversal potential for GABA and blocked by bicuculline (Figure 4C). We also detected optically evoked EPSCs in pyramidal neurons but not interneurons; these events were absent at the reversal potential for glutamate, and they were not blocked by bicuculline.

These oEPSCs were only seen in hSyn-mCherry labeled pyramidal neurons (Figure 4C). Representative recordings of 10 trials from a single postsynaptic DLX2.0-YFP cell are shown in Figure 4D. The first stimulus in a 25 Hz train had a failure rate of 10%, while the fourth stimulus had a failure rate of 40%; these results are aligned with the failure rates for *CALB2* cell activation shown in Figure 4A. The average amplitude oIPSCs in both post-synaptic populations was 3.30 ± 0.4pA (Figure 4E); the latencies, rise times, and durations of oIPSCs (normalized to event peak amplitude) indicated that light stimulation produced monosynaptic GABA release onto the patched cells (Figure 4D).

We observed optical PSCs in both YFP+ interneurons and mCherry+ pyramidal neurons (6/16 DLX2.0-YFP+ cells (37.5%) and 3/14 hSyn-mCherry+ cells (21.4%)). When performing voltage clamp measures, we found that *CALB2*-ChIEF activation evoked inhibitory, but not excitatory, responses in interneurons (6/16 DLX2.0-YFP+ cells (33.3%) exhibited inhibitory responses and 0/16 cells exhibited excitatory responses upon *CALB2*-ChIEF activation; Figure 4F). By contrast, *CALB2*-ChIEF activation evoked inhibitory responses in some pyramidal neurons, and excitatory responses in others (3/14 hSyn-mCherry+ pyramidal cells (21.4%) exhibited inhibitory responses and 2/14 cells (14.3%) exhibited excitatory responses *CALB2*-ChIEF activation; Figure 4F). It is unclear whether this observation of excitatory responses in pyramidal neurons reflected glutamate release from a subset of *CALB+* cells or off-target expression of the CellREADR ChIEF. Notably, our PatchSeq experiments indicated that at least a subset of *CALB2*-targeted cells, particularly those that align with the PAX6 SYT6 transcriptomic subtype, appear to express both vesicular glutamate transporter (*SLC17A7*) and glutamate decarboxylase transcripts (*GAD1* and *GAD2*); Figure 2B)); additionally, a small subset of human MTG excitatory neurons express low levels of *CALB2* (Supplemental Figure 2A)^59^. These results may indicate that neurotransmitter signaling from human *CALB2* neurons may differ from that produced by rodent *Calb2* neurons. We did not observe both excitatory and inhibitory responses to *CALB2*-ChIEF activation in the same pyramidal cell.

### Cell type-specific monitoring of population activity with CellREADR

Last, to investigate CellREADR functionality for imaging cell type population activity, we used binary vectors to express the genetically encoded calcium indicator, GCamp7f, in *CALB2* and *FOXP2* target populations. To establish how calcium signals reported membrane depolarizations and action potentials, we first measured fluorescence signals from individual GCaMP7f+ cells while performing current injections with patch clamp (Figure 5A). In both target populations we observed a linear relationship (R^2^ of the linear fit *CALB2*: 0.84; *FOXP2*: 0.93) between the GCaMP ΔF/F measure and the frequency of evoked action potentials. During current clamp recordings, we also observed both prolonged spontaneous subthreshold membrane depolarizations and spontaneous action potentials in the same cell; these events were reflected by distinct fluorescence signals with different rise times (Tau_3AP_ = 390 ms, Tau_Depolarization_ = 1760 ms, Tau_1AP_ = 202 ms, Figure 5B).

As we had observed frequent spontaneous action potential firing in *CALB2-* and *FOXP2*-targeted neurons under patch clamp (Figure 3), we expected GCaMP imaging would detect population activity in the slice preparation. Indeed, when performing wide field imaging of neocortical slices without stimulating agents, we observed baseline spontaneous activity in both populations (representative images of GCaMP labeling, Figures 5C). Representative time series of ΔF/F signals for *CALB2-* and *FOXP2*-targeted populations are shown in Figure 5D. Qualitatively, spontaneous population activity patterns varied between slices, even across slices that were prepared from the same subject’s tissue. Action potentials (reflected by fast-rising calcium transients of varying amplitudes (as in Figures 5A-B)) were observed in multiple cells from the READR-targeted populations (Figure 5D). We also observed prolonged calcium signals (as in Figure 5B) that occurred simultaneously in many cells (Figure 5D). These latter events may reflect the influence of local neuromodulatory signals or enhanced within-population connectivity across the imaged population. Coordinated activity was more prominent in *FOXP2*-targeted populations than in *CALB2*-targeted populations, as reflected by a cross-wise correlation analysis of calcium signal patterns (Pearson’s rho value for population correlation, 0.639 for the *FOXP2-targeted* cells, 0.402 for the *CALB*-targeted cells; translated into Fisher’s Z values for statistical comparison, p=0.03; Figure 5E).

Differential gene expression analyses of an existing RNAseq database^59^ indicate the expression of nicotinic acetylcholine receptor subunits differs significantly between *CALB2* and *FOXP2* neurons. In particular, cholinergic receptor nicotinic alpha 2 subunit (*CHRNA2)* is expressed in a higher percentage of *CALB2*+ neurons (52.5%) than *FOXP2*+ neurons (1.1%), and at an approximately 1000-fold higher expression level (p-adj = 0), though *CHRNA4*, *CHRNA6*, and *CHRNA7* also showed enrichment in *CALB2*+ over *FOXP2*+ neurons (p-adj = 8.75x10^-7^, 3.22x10^-94^, 1.27x10^-191^, respectively). Of nicotinic acetylcholine receptor subunits, only *CHRNA1* showed enrichment in *FOXP2* neurons (2.8% of FOXP2+ neurons, with approximately 100-fold enrichment over *CALB2* neurons, p-adj = 6x10^-3^). We thus hypothesized that calcium signal responses to nicotinic acetylcholine receptor signaling would differ between these populations. We compared transients from cells in the absence and presence of bath nicotine, and observed a trend towards increased engagement of *CALB2*-targeted cells in population activity (calcium signaling events were detected in 19/100 *CALB2*-targeted cells (19%) under baseline conditions, and in 30/100 cells (30%) following nicotine application; calcium signals were detected in 32/65 *FOXP2*-targeted cells at baseline (49.2%) and in 33/65 cells (50.8%) following nicotine application (n=3 slices for each); *CALB2,* p=0.07 and *FOXP2,* p=0.999 by Fisher exact test, Figure 5G top). In active *CALB2*-targeted cells, nicotine application was found to increase calcium transients relative to baseline conditions (*CALB2*-targeted cells: 2.4 ± 1.1 ΔF/F under baseline conditions, 6.0 ± 2.6 ΔF/F following nicotine application, 2.52 ± 0.9 fold increase; *FOXP*2-targeted cells: 1.6 ± 0.4 ΔF/F under baseline conditions, 1.0 ± 0.4 ΔF/F following nicotine application, 0.94 ± 0.6 fold change; 2-way ANOVA interaction, p=0.045, Figure 5G bottom).

## DISCUSSION

To advance our understanding of the human brain and devise more effective neurotherapies, it is essential to investigate the organization and function of human neural circuits at cell type-resolution. *Ex vivo* human tissues hold significant potential as a platform for studying human neural circuits, but the lack of facile and scalable tools for targeting cell types limits the discovery and translational impacts of this model system. Here we demonstrate that the CellREADR RNA sensor-effector technology enables specific, effective, and programmable recording and manipulation of marker-defined neuronal types in *ex vivo* human brain tissues. Our findings highlight CellREADR’s promise as a genetic tool for human neuroscience and therapeutic applications.

Specificity is of foremost importance for cell type-targeting technologies. While we found that *CALB2* and *FOXP2* sensors exceeded 75% specificity, based on immunohistochemical validations (Figure 1), these measures may be underestimates, as immunohistochemistry is technically challenging in human slice cultures due to factors such as lipofuscin autofluorescence and reagent availability (e.g., many commercial antibodies are not raised against human antigens). Indeed, our single-cell PatchSeq, which assessed comprehensive transcriptional profiles of READR-labeled cells, demonstrated 95% specificity (18/19 cells tested) for the *CALB2* sensor (Figure 2). Our ongoing testing of multiple sensors for *PVALB*, *SST*, and *UNC5B* has yielded promising READRs for other cell types (Figure 1C), but quantitative validation awaits further testing through RNA *in situ* hybridization, antibody staining, or RNA-seq analysis. Future studies will implement large-scale screens of sensor libraries in human cell culture systems and *ex vivo* tissues. Such efforts will facilitate sensor identification and design algorithms and ultimately establish comprehensive RNA sensor libraries for human neuron types.

Payload expression level is also critical to the functionality of a cell-targeting tool. While CellREADR can potentially be delivered as *in vitro* transcribed mRNAs using lipid nanoparticles^73,74^, here we focused on DNA and AAV-based delivery, which may be better-suited for long-term payload expression in brain cells. In our singular vector design, direct coupling of payload translation to the sensor yielded mNeonGreen fluorescence in live tissues 6-9 days after virus application. The binary vector design substantially enhanced payload expression through tTA-TRE transcriptional amplification, achieving live mNeonGreen labeling within 3 days of virus application. Also, the binary vector system expressed fluorescent proteins, calcium indicators and opsins at levels that enabled targeted patch clamp (Figures 2 and 3), optogenetic manipulation (Figure 4), and calcium imaging (Figure 5) of human cell types. As CellREADR is orthogonal to and complementary with transcriptional enhancer-based tools, we were able to use it together with an interneuron class-specific enhancer, DLX-2.0, to investigate human interneuron synaptic connectivity (Figure 4). Our optogenetic data extend from previous studies^19,37,39^ to provide the first proof-of-principle for cell type-specific optical interrogation of synaptic connectivity in human tissues. Likewise, our CellREADR calcium imaging experiments (Figure 5) also stand as the first demonstration of cell type-specific calcium imaging in human microcircuits^39^.

Capsid (i.e., serotype) selection is prerequisite to effective viral delivery and expression of the genetic payload in the target population, and should be validated from the outset of CellREADR applications^25^. The AAV-PHP.eB (as in Lee et al.^33^) and rAAV2-retro (here) appear to have broad tropism for human neuron classes, and thus can be paired with CellREADR for targeting neuronal types. In addition, the promoter used to drive READR expression must be active in the targeted cells. Here we used a human synapsin promoter^75^, which appears to be broadly active in mammalian neurons. Prior to pairing with CellREADR, candidate promoters should likewise be validated in populations that include the target cell type.

To date, transcriptional enhancers are the best-described tools for targeting NHP and human brain cell populations. For example, the DLX2.0 enhancer^21^ provides cross-species (NHP, human, as well as mouse) access to a broad set of cortical GABAergic interneurons^33^, and the mouse *Scn1a* enhancer targets the parvalbumin+ subclass in several mammalian species, including humans^19^. Additional interneuron subclass-specific enhancers have recently been identified through screening and validation in mice, and some of these may be of utility for targeting human interneurons^76^. With advances in single cell epigenome analysis and large-scale enhancer discovery, this approach will continue to yield useful cell type targeting tools, which could also be combined with CellREADR sensors in the same vector for increased specificity. On the other hand, as enhancer screening and validation are typically carried out in mice, the degree to which they are conserved and thus effective in human cell “homologs” remains to be established, though some encouraging results have emerged^19,21,22,76^. Also, recent findings indicate the human brain contains transcriptomic cell types that are unique from those in other mammals^9,46,47^. Studying these human-unique cell types and their related circuits may require dedicated tools; in this context, CellREADR RNA sensors may be better suited to directly target these cells’ specific RNA markers. Alternatively, sensors can be selected for conserved RNA targets, thereby enabling utilization across model organisms. The *CALB2* and *FOXP2* sensors used here were designed specifically for human sequences, but future sensor designs may prioritize such cross-species functionality.

In summary, our results demonstrate that the CellREADR RNA sensor-effector technology can achieve specific, effective, and versatile monitoring and manipulation of human neuron types in *ex vivo* brain tissues. Significant efforts, including large-scale sensor screens in cell cultures and organotypic brain preparations, will be necessary to establish comprehensive sensor tools for human brain cell types. Future efforts will also be directed to refining the specificity of cell type access through the intersectional targeting of 2 or more markers and devising multiplex strategies for differential recording and manipulation of multiple cell types in the same preparation. Other applications may pair CellREADR with a range of tool sets for circuit breaking and cellular control, including neurotransmitter reporters, voltage indicators, retrograde transsynaptic tracers and sonogenetic effectors. Synergistic with and complementary to capsid engineering and enhancer-based approaches, CellREADR will elevate the human *ex vivo* tissue system as a foundational platform and provide a translational bridge towards cell type and circuit specific therapeutics in the human brain.

## Supporting information

All Sup Figures and Tables

## RESOURCE AVAILABILITY

### Lead contact

Further information and requests for resources and reagents should be directed to and will be fulfilled by the lead contact, Derek Southwell

### Materials availability

All unique materials generated for this study are available from the lead contact.

### Data and code availability

Any additional information required to reanalyze the data reported in this paper is available from the lead contact upon request.

## ACKNOWLEDGMENTS

This study was supported in part by NIH grants 1DP2MH140149 (D.G.S.), 1DP1MH129954-01 (Z.J.H.), and 1K08NS133292 (J.B.R). J.B.R. was supported by the NINDS K12 Child Neurologist Career Development Program. D.G.S was supported by the NINDS K12 Neurosurgery Research Career Development Program, The Klingenstein-Simons Fellowship Award in Neuroscience, The Whitehall Foundation, and the Ruth K. Broad Foundation for Biomedical Research. The content is solely the responsibility of the authors and does not necessarily represent the official views of the National Institutes of Health. We thank the Duke Light Microscopy Core Facility and the Duke Sequencing and Genomics Core for their support in data acquisition and analysis. We are grateful to the patients who donated surgical tissue specimens for this research.

## AUTHOR CONTRIBUTIONS

Concept and methodology: E.A.M., D.G.S., Z.J.H., J.B.R., Y.Q.

Data acquisition and analysis: E.A.M., J.B.R., S.Z., P.T., M.M.

Resources: D.G.S., M.L.V. (surgical tissue specimens); Y.Q., S.Z., Z.J.H. (CellREADR viruses)

Writing: D.G.S., E.A.M., Z.J.H., J.B.R.

Funding: D.G.S., Z.J.H.

## SUPPLEMENTAL INFORMATION

Document S1. Figures S1–S5 and Table S1 & S2

## STAR METHODS

## KEY RESOURCES TABLE

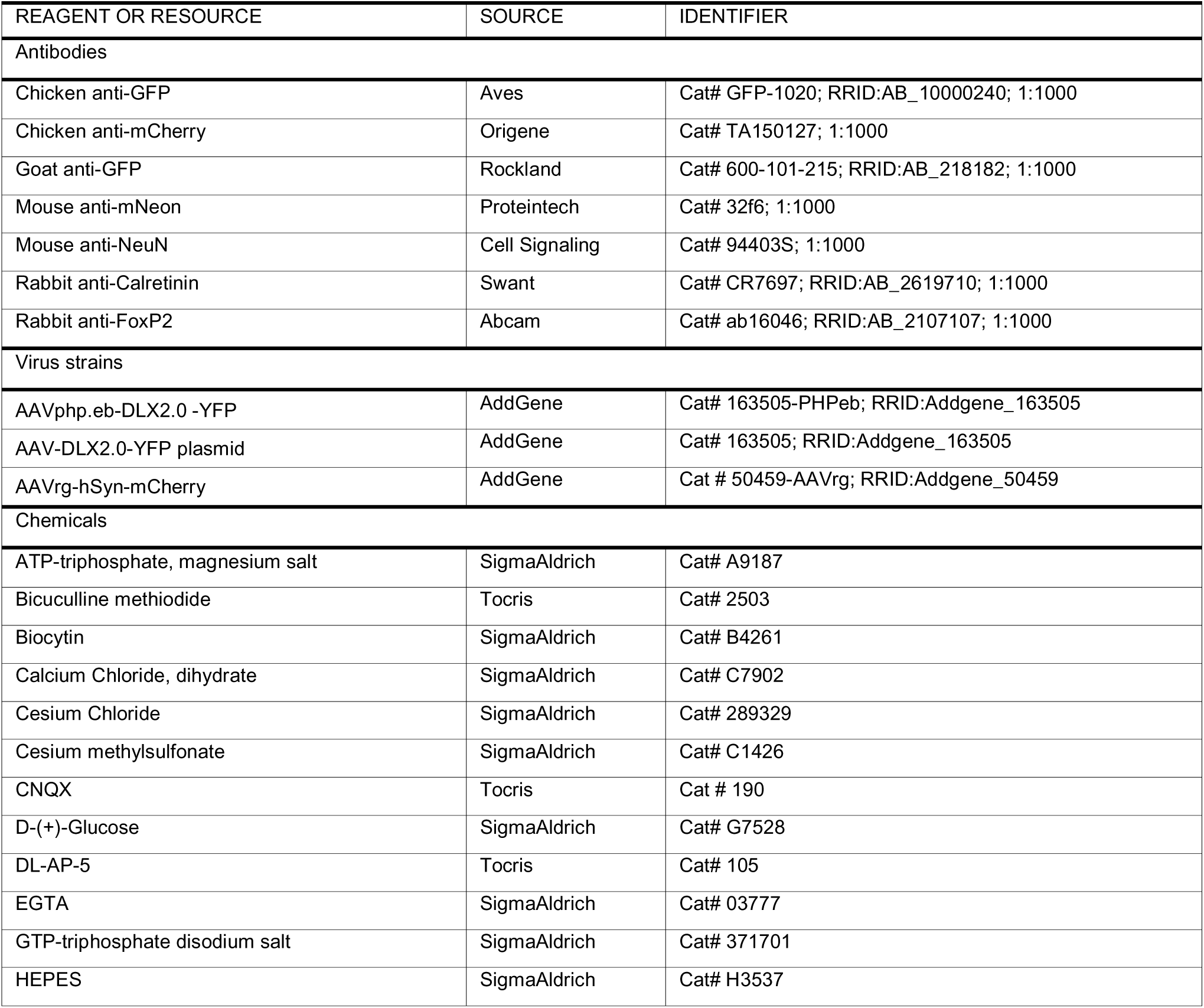
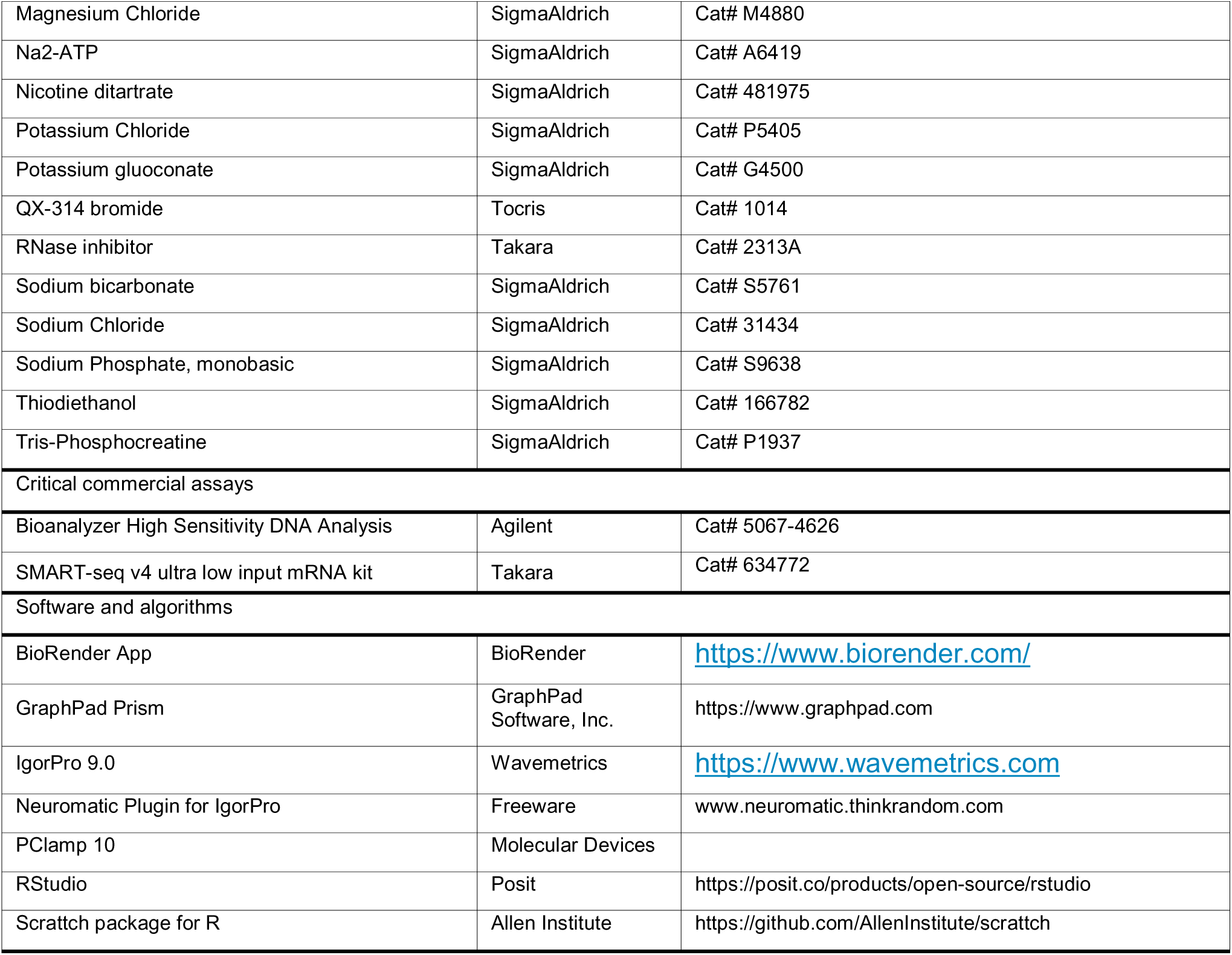

## EXPERIMENTAL MODEL AND STUDY PARTICIPANT DETAILS

### Human tissue procurement

Human cortical tissue was donated by patients who underwent neurosurgical treatments for epilepsy. Non-diseased tissue, which was surgically removed to access areas of disease pathology, was used for this study. All donors provided informed consent under an IRB Protocol (00103019) at Duke University. Biological sex was recorded, but gender identity was not collected. Donor demographics are listed in Table 1.

## METHOD DETAILS

### Plasmids

CellREADR sensors were generated using gBlock synthesis (Integrated DNA Technologies). Vector backbones were linearized using restriction digestion and DNA fragment inserts were generated using PCR or gBlock synthesis (Integrated DNA Technologies).

### CellREADR sensor RNA (sesRNA) design

The sensors used in this study were designed as previously described^48^. In brief, sensors designed for both coding sequences and 5’ untranslated regions of each target gene, depending on the gene length. Synthesized sensor sequences were cloned into CAG-BFP-sesRNA-GFP constructs. We also synthesized the target gene (coding sequence and 5’UTR) using gBlock and cloned them into expression plasmids with CAG promoter. We tested the efficiency and specificity of some sensors in HEK cells using methods we have previously described^48^, and, when performed, used those with the highest specificity for further study. Plasmids contained sequences to promote stability and intended cleavage of the transduced product (Clip-F, WPRE, W3SL)^78^ .

### Viruses

CellREADR viruses were designed and prepared using methods we have previously described^48^. In brief, for production of READR and reporter viruses, HEK293T cells were transfected with READR or Reporter plasmids, AAV serotype plasmids, and pHelper using PEI MAX (Polysciences, 24765). Seventy-two hours after transfection, the cells were collected in cell culture medium, and centrifuged at 4,000 rpm for 15 min. The supernatant was discarded, the pellet was resuspended in cell lysis buffer, frozen, and thawed three times using a dry ice/ethanol bath. Cell lysate was centrifuged at 4,000 rpm for 20 min.

Contaminating DNA in the supernatant was removed by benzonase treatment. The crude viral preparation was loaded onto an iodixanol density gradient and spun at 60,000 rpm for 90 min using a Beckman Ti70 rotor. After the centrifugation, 3–4 ml of crude viral preparation was collected from the 40–60% layer with an 18-gauge needle attached to a 10-ml syringe. The viral crude solution was concentrated using Amicon Ultra-15 centrifugal filters (100 kDa), washed with 8 ml PBS once, and concentrated to an appropriate volume.) AAVrg-DLX2.0-YFP (Addgene #163505) and AAVrg-hSyn-mCherry (Addgene #50459) were used to target interneurons or pyramidal neurons, respectively.

### Human organotypic sample preparation and culture

Organotypic slices from surgical samples were prepared according to published methods^31,32^. Dissection solution contained (in mM): 75 sucrose, 87 NaCl, 25 glucose, 26 NaHCO_3_, 2.5 KCl, 1.2 NaH_2_PO_4_, 10 MgCl_2_, 0.5 CaCl_2_, bubbled with 95/5% O_2_/CO_2_ (pH 7.4). Cortex was carefully dissected under ice cold (∼4C) oxygenated dissection solution to remove the pia. Single gyri were blocked for slicing and cut at 350μm thickness on a vibratome. Slices were rinsed in Hank’s Buffered Saline and plated on PTFE membranes (Millipore, cat. # PICMORG50) in 6 well plates. Tissue was cultured in a conditioned media^31^. Conditioned media also contained antibiotic/antimycotic (1X, Gibco by ThermoFisher) for the first 7 days *in vitro*, and cyclosporine (5 μg/mL) for the entire culture period. HEPES (20mM) was added to the media for the first hour after slice preparation. Slices were cultured in an incubator at 35 Celsius with 5% CO_2_ and 5% humidity. Virus-mediated expression of fluorescent reporters was monitored beginning at 3 DIV until they were used for immunohistochemistry (DIV 3-16), patch clamp electrophysiology (DIV 4-12), or calcium imaging experiments (DIV 4-7).

### Virus application

On the day of tissue donation, after slicing and plating the tissue on semi-permeable membranes and incubating slices in HEPES containing media, viruses were directly applied to the surface of the slice. Viruses were diluted in media (CellREADR virus, 1:8 dilution; TRE-effector viruses, 1:6 dilution; AAVrg-DLX2.0-YFP and AAVrg-hSyn-mCherry 1:10 dilution), and 7µL of the virus solution was applied to slice grey matter. Plates were carefully returned to the incubator and left undisturbed overnight.

### Immunohistochemistry

After 7 of incubation, or upon completion of patch clamp recordings, cortical slices were fixed overnight in 4% PFA, then rinsed in PBS and stored for 1-7 days in PBS with azide. For biocytin recovery or colocalization IHC, tissue was permeabilized in PBS with 10% TritonX100, 5% NGS, and 100µM glycine 2 hours at room temperature on a shaker plate.

Primary antibodies were incubated 48 hours at 4C; secondary antibodies were incubated for 24 hours at 4C. Biocytin was developed with either Alexa647 conjugated streptavidin, or with a silver stain. Large overview images of native mNeon fluorescence were made on a Leica SP8 upright confocal microscope, or a Keyence BZ-X800 with a 10X or 20X objective.

The following antibodies were used for immunohistochemical labeling of mNeon, mCherry, FoxP2, Calretinin and NeuN: mouse anti-mNeon Ab 1:500, Proteintech 32f6; chicken anti-mCherry Ab 1:500, Origene TA150127; rabbit anti-FoxP2 Ab 1:500, Abcam ab16046; rabbit anti-Calretinin Ab 1:500, Swant CR7697; mouse anti-NeuN Ab 1:500, Cell Signaling #94403S. Secondary antibodies were Alexa488, Alexa594, or Alexa647 against the appropriate species (Invitrogen). Slices were mounted using a medium containing DAPI.

Some slices were selected for chemical clearing to enhance imaging deeper into the slice and reduce lipofuscin autofluorescence. These slices were incubated in thiodiethanol (TDE) solution of 30% for 35 minutes, 60% for 35 minutes, then 75% for 10 minutes. Slices were mounted in a 300µm thick spacer ring (SunJin Labs, iSpacer, IS307), then the remaining volume was filled with 75% TDE before coverslip placement.

### Image analysis for colocalization & cell location

Immunohistochemistry measurements of CellREADR specificity were performed by immunostaining against either CALB2 or FOXP2 protein and CellREADR mNeonGreen. Maximum projection images of small z-stacks (40-60 μm total depth) were used for manual counts of mNeon-labeled cells and CALB2 or FOXP2 using the CellCounter Plugin in FIJI. Colocalization was expressed as the percentage of mNeonGreen+ cells that also expressed the marker protein of interest (CALB2 or FOXP2).

### Electrophysiology

Slices were transferred to a heated recording bath (32-34C) on an Olympus BX-50 upright microscope. Recording solution contained (in mM): 118 NaCl, 3 KCl, 25 NaHCO_3_, 1 NaH_2_PO_4_, 1 MgCl_2_, 1.5 CaCl_2_, 30 glucose fully saturated with 95/5% O_2_/CO_2_. Borosilicate patch pipettes were pulled with a resistance of 3.5-5.5 MΩ, and filled with an internal solution containing (in mM): 134 K-gluconate, 10 HEPES, 4 ATP-triphosphate Mg salt, 3 GTP-triphosphate Na salt, 14 Phosphocreatine, 6 KCl, 4 NaCl, pH adjusted to 7.4 with KOH for current clamp recordings For voltage clamp recordings (optogenetic stimulation of PSCs), the internal solution continued (in mM): 100 Cs-MeSO3, 40 CsCl, 4 MgCl2, 0.1 Na2-GTP, 4 Na2-ATP, 10 Tris-Phosphocreatine, 10 HEPES, and 4 QX-314. Biocytin (0.5%) was added to the internal solution to allow for morphological identification after recording. Patched cells were held in whole-cell mode for a minimum of 10 minutes to ensure complete filling with biocytin.

Labeled cells were targeted for patch clamp under a 60X objective. Cells were sampled from across cortical layers, in manners that evenly reflected where the *CALB2* and *FOXP2* CellREADR constructs were expressed (Figure 1). Recordings were performed using an Axopatch 700B amplifier (Molecular Devices), and data were digitized (Digidata 1550B) and captured with pClamp 10 (Molecular Devices). Membrane voltage was recorded at 100k Hz and low-pass filtered at 10k Hz. Liquid junction potential was not corrected. Pipette capacitance and series resistance were compensated at the start of each recording and checked periodically for stability of the recording configuration. Intrinsic membrane properties (input resistance, capacitance) were measured with a -10 pA current step. Rheobase was measured with a ramp current of 1s duration and 100-300 pA final amplitude. Input-output curves were generated from a series of current steps starting at -100 pA and increasing in 10 pA increments until a maximum firing rate was elicited. Sag ratio was calculated from the -100pA step, and was the difference between the negative peak deflection from baseline and the steady-state voltage normalized to the peak deflection^63^. In a subset of cells glutamatergic blockers (20 µM CNQX + 50 µM AP-5) were bath applied to block excitatory synaptic inputs. Data were analyzed using NeuroMatic and custom scripts in Igor Pro (Wavemetrics).

Optical stimulation experiments were performed by expressing either AAVrg-DLX2.0-YFP, AAV9-hSyn-GFP or AAV9-hSyn-mCherry in putative postsynaptic cells, and CALB2-ses1-tTA and TRE-ChIEF-YFP or TRE-ChIEF-tdTom CellREADR constructs in putative pre-synaptic populations. Cells that 1) expressed only the DLX2.0-YFP or hSyn-mCherry, and 2) were situated in areas of CellREADR labeling, were targeted for voltage clamp recordings. Blue light pulses (625 nm, 5x 2ms-long pulses at either 10Hz or 50Hz) illuminating the whole field of view were triggered for 10 sweeps while recording membrane currents at -30 mV (V_rev_ GABA) and at 0 mV (V_rev_ Glu). If an optical response was detected in the average of the 10 sweeps, then a GABAergic blocker (50 µM bicuculline) was bath-applied to the slice, then stimulation was repeated to identify the neurotransmitter that produced the PSC.

### Processing of cellular nuclei for PatchSeq

PatchSeq electrophysiology was performed using an internal solution that included (in mM): 110 K-gluconate, 4 KCl, 10 HEPES, 1 ATP-Mg, 0.3 GTP-Na,10 Na-phosphocreatine, 0.2 EGTA, 20 μg/mL glycogen, 0.5% biocytin, and 0.5 U/μL RNase inhibitor (Takara). The internal solution was prepared with RNase-free water and stored in daily aliquots at -80 C until use. Equipment, materials preparation and nuclei extraction followed previously published protocols^79^. Clean nucleus extraction was confirmed visually with both DIC and fluorescent images. Nuclei were stored at -80C in 10μL of lysis buffer with RNase inhibitor (40 U/µL) and then processed for sequencing in batches (5-12 nuclei/batch). Nuclear RNA was processed into cDNA libraries using the SMART-Seq v4 Ultra Low Input kit (Takara) using 19 PCR cycles. Quality checks of the resulting cDNA were made on a Bioanalyzer (Agilent, High Sensitivity DNA kit). Nuclei that yielded >200 pg/µL cDNA were sequenced. Borderline cDNA (concentrations just below 200 pg/µL, and fragment sizes >500 bp) were re-amplified with 3 rounds of PCR, rechecked for cDNA quality, and included in the sequencing set if reamplification increased cDNA concentration above threshold.

### RNA sequencing and transcriptomic analysis

cDNA sequencing was performed at the Duke Sequencing and Genomics Technologies core facility on an Illumina NovaSeq 6000 flow cell with a targeted sequencing depth of approximately 50 million reads per sample. Data were aligned to the human genome (GRCh38, annotations from ENSEMBL v106). Raw count matrices from aligned data were analyzed using the Allen Institute scrattch.hicat (version 1.0.0) package (https://github.com/AllenInstitute/scrattch.hicat) in R (2022.07.1 Build 554). Normalized counts were calculated as log2(counts-per-million reads per cell) for downstream analysis. Heatmaps for select genes were generated using the gplots package (version 3.1.3) in R.

Predicted transcriptomic cell type annotations were derived from a reference dataset of 15,928 nuclei from the human middle temporal gyrus (MTG) that underwent SMART-seq single nucleus RNA sequencing^59^. Annotation predictions were generated using scrattch.hicat (version 1.0.0) according to a standard data analysis pipeline (https://taxonomy.shinyapps.io/scrattch_tutorial/#section-overview). The reference MTG dataset was limited to interneuron clusters (class = GABAergic); cell class and cluster (i.e. Inh L1-3 PAX6 SYT6) were used to generate reference cluster names. The reference count matrix and the PatchSeq test count matrix were both normalized (log2(counts-per-million(raw matrix) +1)) and key cluster specific genes were selected using the “select_markers” function in scrattch.hicat. Cluster predictions for PatchSeq cells were then generated using the “map_sampling” function in scrattch.hicat (markers.perc=0.8, iter=100).

To investigate differential gene expression between *CALB2*+ and *FOXP2*+ neurons we utilized an existing single-nucleus RNA sequencing atlas from human MTG^59^ . The atlas dataset was reconstituted in Seurat via CreateSeuratObject, and then the raw count matrix was normalized and scaled via SCTransform with default parameters. The human MTG Seurat object was further subsetted for cells with non-zero expression of *CALB2* or *FOXP2*, yielding *CALB2*+ GABAergic neurons and *FOXP2*+ Glutamatergic neurons. Differential gene expression was then analyzed between these two populations via FindMarkers(object, ident.1 = “GABAergic, ident.2 = “Glutamatergic”, only.pos = FALSE), yielding log2-fold changes in expression, percent gene expression in ident.1 and ident.2, and a p-value adjusted for multiple comparisons where p-adj <0.05 was considered significant differential expression. The list of differentially expressed genes was then manually investigated for significantly differentially expressed genes of interest related to neuromodulation (e.g.. nicotinic acetylcholine receptor subunits), for the studies of calcium imaging.

### Morphology

For cortical layer identification, 4 sections from 4 donors were labeled with DAPI and imaged from the pia to the white matter border. The density of nuclei delineated layer boundaries; the distances between each layer were calculated for the 4 samples and averaged to create a “typical” cortical layer map. Our cortical layer distances were as follows: L1-L2 boundary, 221.8 μm, L2-L3 boundary, 383.9 μm, L3-L4 boundary, 812.2 μm, L4-L5 boundary, 1677.6 μm, L5-L6 boundary, 2206.9 μm, L6-WM boundary, 3549.5 μm. This is a slightly larger average thickness than reported elsewhere (2963μm)^38^. When a biocytin-filled cell was identified, the distance from the pia to the cell soma was measured, as was the underlying Layer 6-white matter boundary (these were measured along a line perpendicular to the overlying pia). Soma position was then calculated as a percent depth (0% is located at the pial surface and 100% is located at the Layer 6-white matter boundary. This normalized the depth position of the soma across slices. DAPI labeling was used as a secondary criterion to ascertain the soma laminar position.

For morphological tracings, biocytin-filled neurons were imaged using a 40x objective on a Leica SP8 upright confocal microscope (oil immersion, NA = 1.3). Tiled z-stacks containing cell dendrites and axons were captured at 1024x1024 resolution, with a z-step size of 1.0-1.2 μm. Overview images were captured using a 10x objective to provide a reference for cell orientation relative to the pial surface. Images were traced using the automatic tracing plugin in Vaa3D^80^ and converted to .SWC format. The automatic tracing function sometimes produced erroneous branching points, and so all traces were inspected and manually corrected using either the SNT plugin in FIJI, or neuTube, a plugin for Vaa3D. To orient the traces relative to the pial surface, corrected .SWC files were aligned in the SNT Viewer using the 10X overview images to position all cells vertically in the cortical column.

### Calcium Imaging

Four to 7 days after virus application, slices with robust GCaMP7f fluorescence were selected for experiments. Slices were placed in a bath chamber at 32 C under a 10x objective and a high-power LED (Thor Labs) at 488 nm. Videos of 60-90 s duration were captured at a frame rate of 100 ms with a Retiga Electro 1.4 M pixel CCD camera (Teledyne Photometrics) with no binning. Imaging was performed in supragranular cortex, where the greatest density of cellular labeling was observed with both the *CALB2* and *FOXP2* CellREADRs. FIJI was used to check the standard deviation projection of the video for spontaneous activity. If activity was detected, several videos were then captured in the same region, then a different area of the slice was imaged. Between 4 and 10 videos were captured from each field of view over the course of 10-15 minutes. Nicotine (300 µM) was bath-applied to a subset of slices, then videos were recaptured 5 minutes later from the same field of view. After allowing nicotine to wash out for 15 minutes, baseline imaging was then performed at a new location.

One representative video of each field of view was selected if it contained a relatively high density of labeled cells and exhibited minimal motion artifacts. Data were analyzed with ImageJ (FIJI) and custom scripts in Igor Pro. If necessary, motion artifacts were corrected with the Image Stabilizer plugin.

Maximum projection images were used to select all virally neurons in the field of view for analysis (non-active cells were also analyzed). ROIs were assigned to each neuron, and intensity values over time were extracted for each ROI. Calcium signals were background subtracted and normalized to a baseline period that exhibited no activity. Co-activation was calculated by a pairwise cross-correlation (Pearson’s) of all the ROIs in the field of view. Pearson’s correlation coefficients were converted to Fisher’s z-values for comparison across slices and between groups. Activity during the nicotine application was normalized to baseline activity in standard recording solution.

## QUANTIFICATION AND STATISTICAL ANALYSIS

All data values in the text are the mean +/- SEM; error bars in the figures indicated the SEM. Statistical tests were performed in GraphPad Prism. Data were checked for normality; normally distributed data were compared with t-test or ANOVA. Non-normal data distributions were compared with the Mann-Whitney U-test. Differences in proportions were tested with the Fisher’s exact test (small sample size) or Chi-squared test.

## ADDITIONAL RESOURCES

Creative Commons license for figure Panels created with the BioRender app:

Graphical Abstract: Created in BioRender. Matthews, E. (2025) https://BioRender.com/0pgjkzm

Figure 2, Supplemental Figure 2, & Supplemental Figure 5: Created in BioRender. Matthews, E. (2025) https://BioRender.com/qhb53fv

