## Supplementary material for "RNA-programmable cell type monitoring and manipulation in the human cortex with CellREADR": All Sup Figures and Tables

### **Supplemental Figure 1**

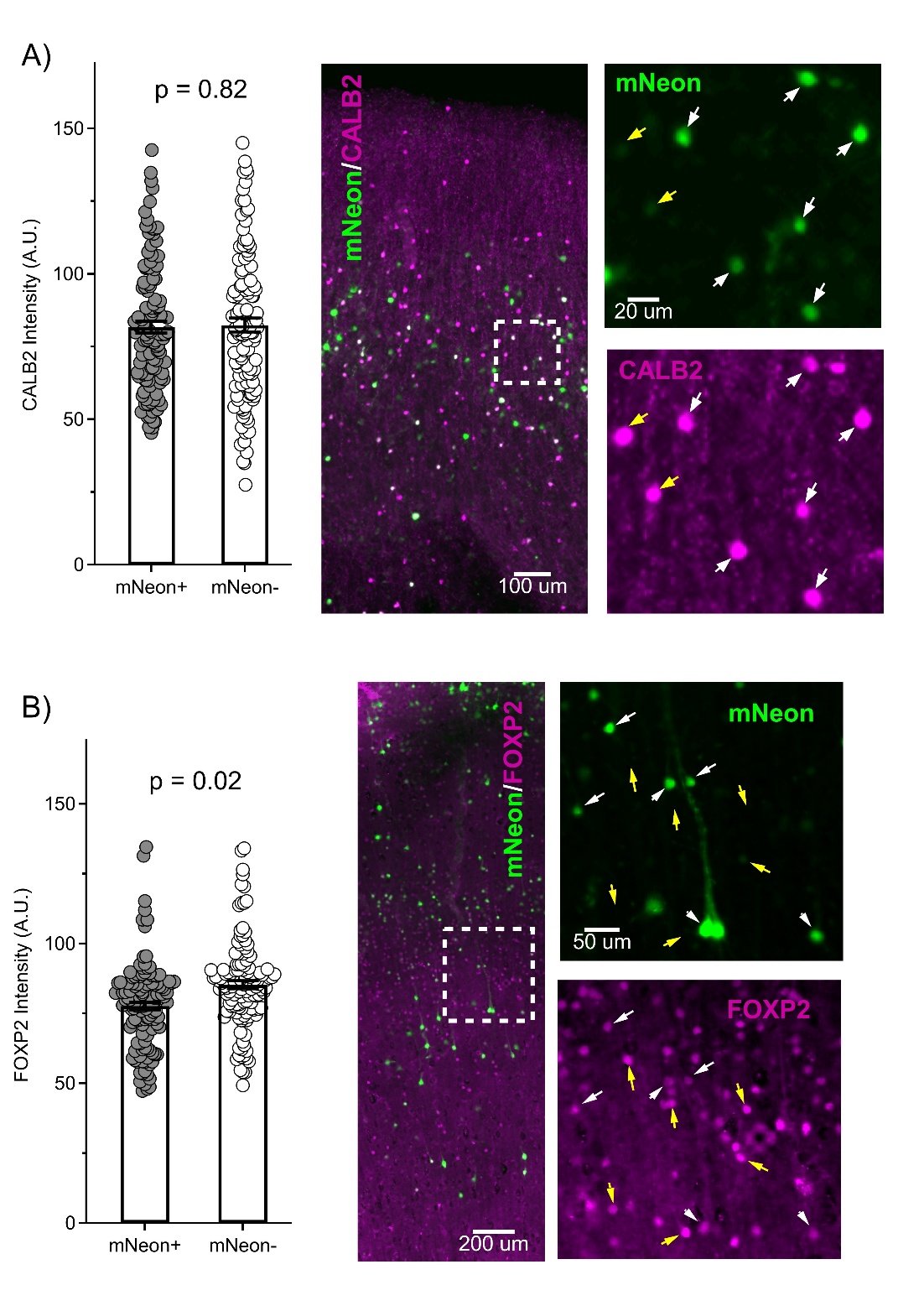

**Supplemental Figure 1: CellREADR does not strongly perturb protein expression of the target RNA**

(A) CALB2 immunostaining signal intensity (purple) was compared between *CALB2* cells that had been successfully targeted by a binary *CALB2* CellREADR (with mNeonGreen reporter, green; designated by white arrows) and cells that were not targeted (yellow arrows).  CALB2 signal intensity did not differ between READR-targeted cells and untargeted cells. Scale bars: 100 µm, 20 µm.

(B) FOXP2 immunostaining signal intensity (purple), compared between cells targeted by a binary *FOXP2* CellREADR (mNeonGreen reporter, green; designated by white arrows) and cells that were not targeted (yellow arrows).  FOXP2 signal intensity was slightly reduced in READR-targeted cells. Scale bars: 200 µm, 50 µm.

### **Supplemental Figure 2**

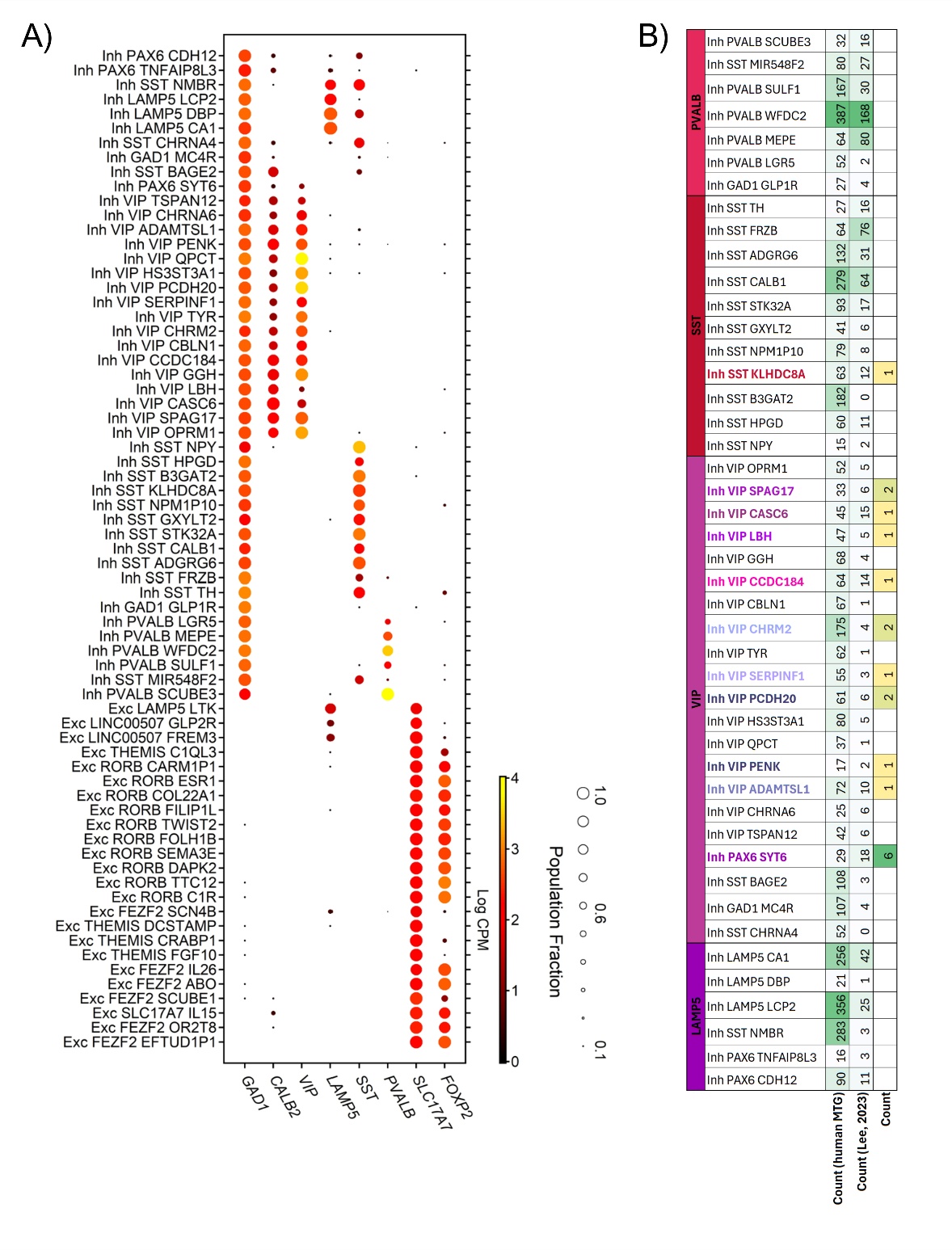

**Supplemental Figure 2: Human MTG transcriptomic data referenced in current study**

(A) Reference gene expression profiles of human MTG neurons constructed by the Allen Institute^1^. *FOXP2* expression is nearly restricted to the excitatory neuron class and expressed widely across excitatory neuron types. *CALB2* is expressed primarily in GABAergic neurons, and in low levels in a few types of excitatory neurons.

(B) CellREADR programmability enables the restricted targeting of an interneuron subclass. Counts of interneuron transcriptomic types identified in a reference atlas obtained primarily with acute post-mortem tissues (left column, human MTG)^1^. Transcriptomic types characterized in a previous PatchSeq study, which utilized the DLX2.0-YFP enhancer virus to target the interneuron class (some cells were also accessed without viral labeling; middle)^2^. Counts of *CALB2* READR-targeted cells that were assessed by RNAseq in the current pilot study; note utility of CellREADR for systematic access to specific populations.

### **Supplemental Figure 3**

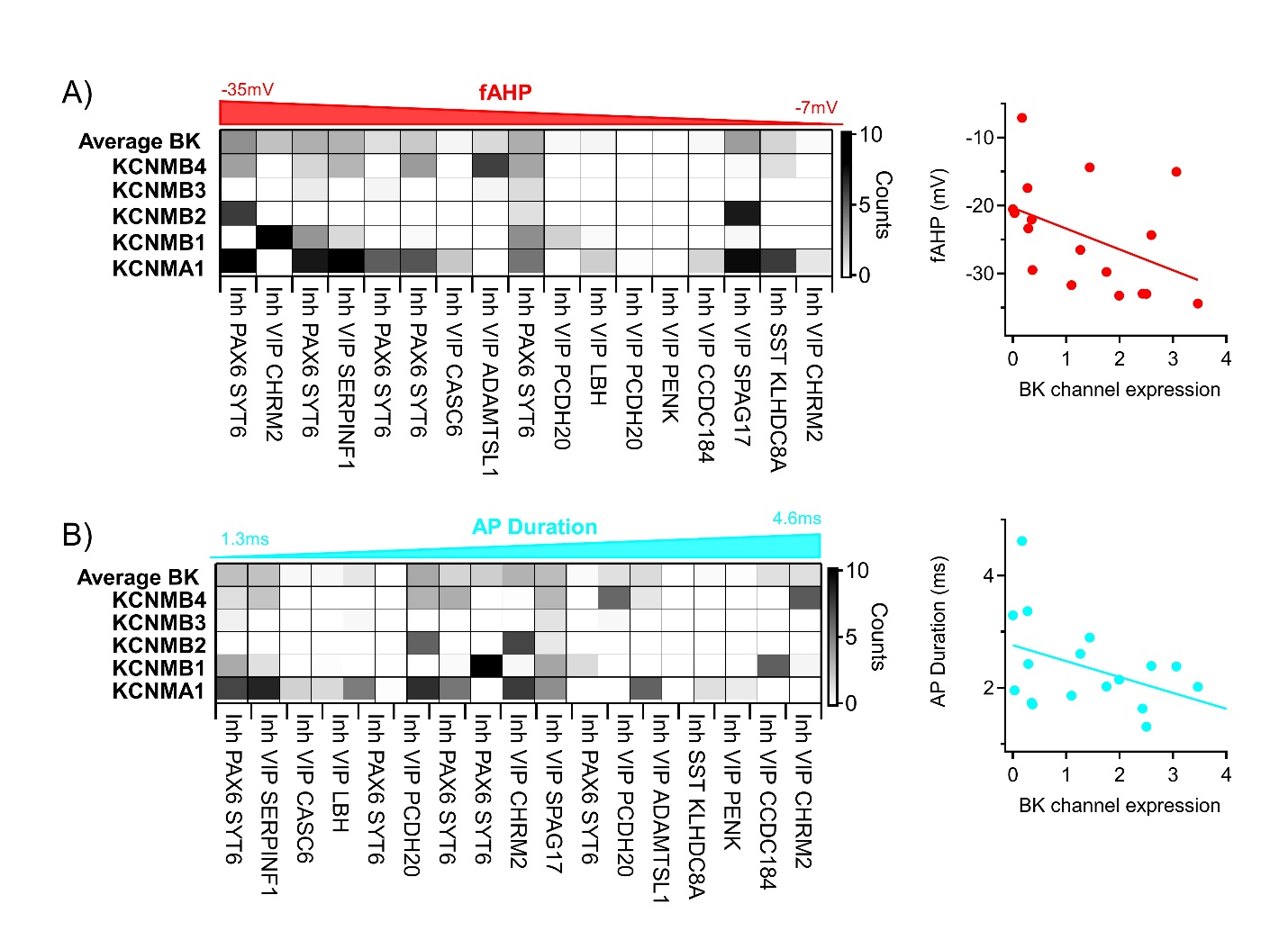

**Supplemental Figure 3: Correlation between gene expression and action potential properties in cells targeted with CALB2 CellREADR**

(A) RNA expression counts for the BK channel alpha (*KCNMA1*) and beta subunits (*KCNMB1-4*) are shown for each cell that completed PatchSeq, with single-cell data sorted from left-to-right by decreasing amplitude of the fast afterhyperpolarization (fAHP) following single action potentials. As expected, the average expression of all BK channel subunits correlates with fAHP amplitude, especially with the pore-forming alpha subunit. The plot at right shows the relationship between average BK channel expression and the fAHP amplitude (R^2^ = 0.19).

(B) RNA expression counts for the BK channel subunits, sorted by increasing action potential duration. As for the fAHP, the expression of BK channel subunits correlates with the AP duration. Plot at right shows the relationship between average BK channel expression and the AP duration (R^2^ = 0.17).

### **Supplemental Figure 4**

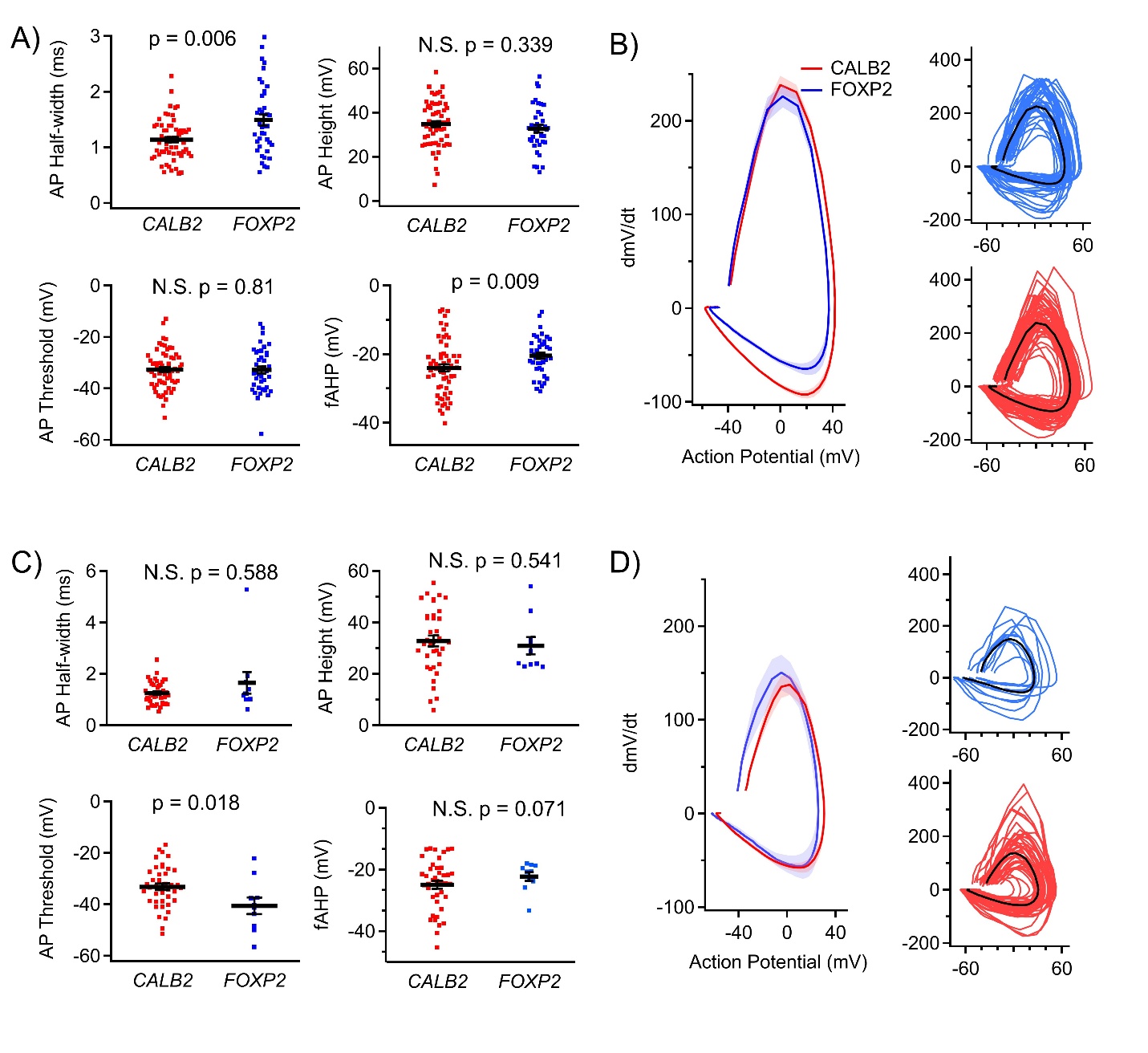

**Supplemental Figure 4: Additional measures of evoked and spontaneous action potentials**

(A) Action potentials elicited by a 500 ms current step at 2X the rheobase current were measured.  Afterhyperpolarization (AHP), peak, half-widths and durations were compared between *CALB2-* (n=60 cells) and *FOXP2-*targeted (n=43 cells) neurons (Mann-Whitney U-test for non-normal distributions, t-test for normal distributions).

(B)  Action potential phase plots for the grand average (left) and individual cells (right) are shown for the *CALB2*- and *FOXP2*-targeted populations. The repolarization phase and fast afterhyperpolarization features shown in (A) are emphasized in the grand average here and indicate that *CALB2*-targeted cells exhibit phenotypes consistent with interneurons.

(C) Spontaneous action potentials recorded at rest (during a 3-minute period where zero holding current was injected) were also analyzed and compared between the *CALB2*- and *FOXP2*-targeted populations (n=42 and n=12, respectively). Differences in the fast AHP & AP half-width, as observed above, were not present in these subsets of spontaneously active cells; however, action potential thresholds were found to differ (Mann-Whitney U test).

(D) Action potential phase plots for the grand average (left) and individual cells (right).

### **Supplemental Figure 5**

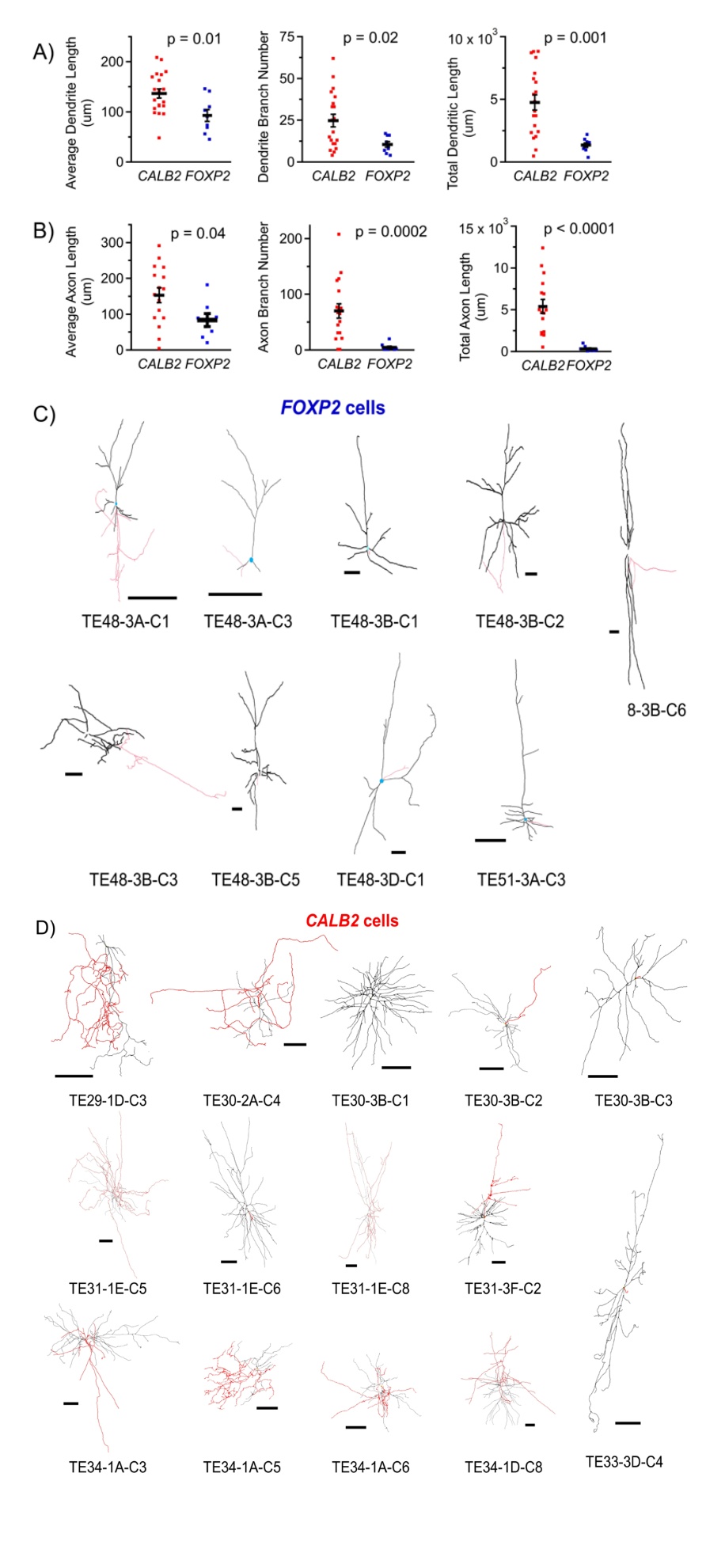

**Supplemental Figure 5: Morphometrics of CALB2- and FOXP2-targeted neurons**

(A) A subset of neurons was recovered for morphological reconstructions and analysis following patch-clamp recordings. Dendritic average lengths (left), branch numbers (middle), and total length (right) were larger in *CALB2*-targeted cells, than in *FOXP2*-targeted cells (Mann-Whitney U-test for non-normal distributions, t-test for normal distributions).

(B) Properties of recovered axons were also measured, when cellular morphology was sufficiently recovered. Often only a short segment of axon was recovered from *FOXP2*-targeted cells. Axons of *CALB2*-targeted cells were more frequently intact and of greater average branch length, branch number, and greater total length than those recovered from *FOXP2-*targeted cells (Mann-Whitney U-test for non-normal distributions, t-test for normal distributions).

(C) Morphologies of a sample of 7 *FOXP2*-targeted cells are shown, with axons in red, dendrites in black, and soma in cyan. Scale bar: 200 µm.

(D) Morphologies of a sample of 14 *CALB2*-targeted cells, which were not analyzed by PatchSeq (see Figure 2).  Axons, dendrites, and soma are labeled as in C. Scale bar: 100 µm.

### **Table S1. Intrinsic physiological properties measured at different periods of slice culture**

| *CALB2*-targeted | Early (DIV 4-7) | Late (DIV 8-12) | p-value |
| --- | --- | --- | --- |
| N (cells) | 28 | 32 |  |
| V_rest_ (mV) | -56.8 ± 1.4 | -53.6 ± 1.5 | 0.121 |
| C_m_ (pF) | 37.9 ± 5.7 | 24.8 ± 3.1 | 0.105 |
| R_input_ (MOhm) | 0.296 ± 0.063 | 0.263 ± 0.052 | 0.863 |
| Max Firing Freq (Hz) | 56.5 ± 4.3 | 51.7 ± 2.7 | 0.338 |
| Gain (Hz/pA) | 0.533 ± 0.064 | 0.586 ± 0.047 | 0.504 |
| Rheobase (pA) | 47.5 ± 10.7 | 27.2 ± 4.7 | 0.077 |
| Spont. Firing Freq (Hz) | 1.91 ± 0.60 | 1.94 ± 0.57 | 0.609 |
| Proportion Spont. Firing (%) | 69.7 | 68 | 0.999 |
| *FOXP2*-targeted | **Early (DIV 4-7)** | **Late (DIV 8-12)** | **p-value** |
| N (cells) | 25 | 18 |  |
| V_rest_ (mV) | -46.5 ± 2.5 | -44.7 ± 3.5 | 0.661 |
| C_m_ (pF) | 92.0 ± 13.3 | 83.5 ± 11.7 | 0.836 |
| R_input_ (MOhm) | 0.077 ± 0.020 | .089 ± 0.033 | 0.741 |
| Max Firing Freq (Hz) | 37.1 ± 4.0 | 34.7 ± 8.5 | 0.777 |
| Gain (Hz/pA) | 0.192 ± 0.28 | 0.170 ± 0.062 | 0.712 |
| Rheobase (pA) | 119.3 ± 17.2 | 98.2 ± 23.9 | 0.465 |
| Spont. Firing Freq (Hz) | 1.80 ± 0.99 | 0.04 ± 0.02 | 0.379 |
| Proportion Spont. Firing (%) | 40.0 | 11.1 | 0.046 |

Data presented in Figure 3 were binned according to when they were collected (DIV 4-7 versus DIV 8-12).  For *CALB2*- and *FOXP2*-targeted cells, intrinsic physiological properties did not differ between early and late periods.  During the late period, a reduced proportion of *FOXP2*-targeted cells exhibited spontaneous firing, though the proportion of CALB2-targeted cells that was firing did not differ between early and late periods.

### **Table S2. Comparison of intrinsic physiological measures to a previous data set**

|  | Current study | Lee et al. (*VIP* only) | p-value |
| --- | --- | --- | --- |
| N (cells) | 60 | 27 |  |
| V_rest_ (mV) | -55.1 ± 1.1 | -61.5 ± 0.6 | 0.001 |
| R_input_ (MOhm) | 0.28 ± 0.04 | 0.31 ± 0.02 | 0.508 |
| Gain (Hz/pA) | 0.56 ± 0.04 | 0.46 ± 0.04 | 0.131 |
| Rheobase (pA) | 51.6 ± 5.7 | 29.8 ± 5.0 | 0.019 |
| AP Threshold (mV) | -32.4 ± 1.0 | -33.7 ± 1.4 | 0.463 |
| AP Half-width (ms) | 1.00 ± 0.05 | 0.84 ± 0.07 | 0.086 |
| Fast AHP (mV) | -24.0 ± 1.0 | -20.6 ± 1.8 | 0.080 |

Patch clamp measures from *CALB2*-targeted cells (Figure 3) were compared to those gathered from human *VIP* subclass interneurons in a prior study, in which the majority of cells were patched in acute slices without viral labeling, with smaller numbers targeted in cultured slices with DLX2.0 virus application^2^. While there were some methodological differences between recording techniques (e.g., patch clamp was performed with synaptic blockers previously; here, it was not) and cells were not necessarily equally sampled, these comparisons generally suggest CellREADR did not grossly alter cellular physiology.
